# SMART-NeuroDx: A Reagent-Free Multianalyte Biosensor Platform with Machine-Learning Readout for Point-of-Care Dementia Screening

**DOI:** 10.64898/2026.07.22.740043

**Authors:** Sudhaunsh Deshpande, Krzysztof Pawlak, Moulya Prathap, Cara Evans, Theresa Al Alam, Anu Mary Joy, Ajith Mohan Arjun, Sanjiv Sharma

**Affiliations:** David Price Evans Global Health and Infectious Diseases Group, Pharmacology & Therapeutics, Institute of Systems, Molecular and Integrative Biology, University of Liverpool, Crown Street, Liverpool L69 7BE, United Kingdom; Materials Innovation Factory, University of Liverpool, 51 Oxford Street, Liverpool L7 3NY

**Keywords:** Molecularly Imprinted Polymers (MIPs), Point-of-Care Biosensors, Machine Learning, Differential Dementia Screening, Alzheimer’s Disease, Frontotemporal Dementia

## Abstract

Differentiating overlapping dementia pathologies, such as frontotemporal dementia and Alzheimer’s disease, calls for the simultaneous measurement of several blood biomarkers, yet electrochemical sensors remain predominantly single-target and dependent on labels and sample preparation. Here we report SMART-NeuroDx, a reagent-free electrochemical array that quantifies pTau217, GFAP, pTau181, and NfL directly from unprocessed plasma and serum in under 35 minutes. Four surface-confined redox-active molecularly imprinted polymer and aptamer-MIP recognition matrices are electropolymerized onto a four-working-electrode porous-gold printed-circuit array, using potential-assisted electrostatic gating keyed to each target’s isoelectric point to prevent cross-channel template contamination during synthesis. Label-free Faradaic responses from the redox-active polymer backbone are acquired on a custom battery-powered STM32G4 handheld potentiostat and converted to concentration from five voltammetric features using cross-validated Random Forest and XGBoost regressors. The handheld unit reproduced the baseline fidelity of a commercial benchtop workstation and resolved sub-picogram pTau217 (limit of detection 0.087 pg mL⁻¹) across marker-appropriate dynamic ranges, with inter-chip relative standard deviation at or below 5.34% and coefficients of determination of 0.94 to 0.99 against reference concentrations. Each channel retained selectivity in plasma and serum against competing neurological and inflammatory proteins, with non-specific signal deviation held below 10% by the hydrated PyPEG interfacial shell. The platform establishes reagent-free, simultaneous four-analyte neurodegeneration sensing on a single point-of-care device.

---

Precise measurement of blood-based neurological biomarkers has transformed the detection of Alzheimer’s disease (AD), establishing single-analyte plasma tests like phosphorylated-tau 217 (pTau217) as highly accurate indicators of amyloid pathology entering standard clinical trial and care pathways ^1–4^. Currently, there exist many well-developed electrochemical sensors optimized for single biomarker detection ^5–12^. However, clinical dementia assessment is inherently a differential problem. Because AD and frontotemporal dementia (FTD) overlap significantly in clinical presentation, particularly in younger and atypical presentations an AD-specific marker by design reports only the probability of AD biology rather than the presence of a non-AD dementia ^13–17^. Due to this heterogeneous and overlapping nature of dementia etiologies, a single biomarker is fundamentally insufficient for a definitive differential screening ^13,18,19^.

We have previously shown that multianalyte sensing facilitates a greater distinction between healthy controls (HC) and AD when put together. With the clinical availability of disease-modifying therapies (DMTs), it is now imperative that point-of-care platforms can differentially diagnose between AD, FTD, and other forms of dementia to prioritize patients for confirmatory clinical pathways ^20–25^. This clinical necessity highlights two critical gaps in the current landscape. First, the assays that achieve high-accuracy AD detection are predominantly central-laboratory tests optimized to identify AD biology rather than to resolve AD from non-AD neurodegeneration ^26–28^. Second, these high-fidelity platforms remain heavily concentrated in relatively few, well-resourced centres, whereas dementia prevalence is rising fastest in low- and middle-income countries (LMICs) where cerebrospinal fluid testing, amyloid-PET, and central-laboratory biomarker infrastructure are scarce ^29^. The populations with the greatest and fastest-growing need are thus the least served by the current generation of tests ^30,31^.

While our previous work successfully established single-biomarker AD diagnostics via the detection of hyperphosphorylated Tau181 (pTau181), we missed an important biomarker, pTau217, which head-to-head comparisons have shown to be a superior discriminator of AD pathology ^5,7^. Furthermore, integrating markers of astrocytic reactivity, such as Glial Fibrillary Acidic Protein (GFAP)—and pan-neurodegeneration markers like Neurofilament Light chain (NfL) captures complementary biological axes across the AT(N) framework directly from blood circulation ^32–35^.

To address these gaps simultaneously, we introduce here the SMART-NeuroDx platform (**Figure 1**): a portable, low-cost, and reagent-free electrochemical system designed for the simultaneous measurement of pTau217, pTau181, GFAP, and NfL directly from undiluted plasma in less than 35 minutes. By measuring these distinct pathological axes in a multiplex format, the SMART-NeuroDx platform uniquely provides automated ratiometric measurements—specifically the NfL: pTau217 ratio—to deliver an FTD-facing readout that captures neuroaxonal injury in the relative absence of AD-type tau pathology. This design offers a pragmatic way to flag non-AD neurodegeneration at the point of care while direct, primary FTD biomarkers mature.

**Figure 1.**
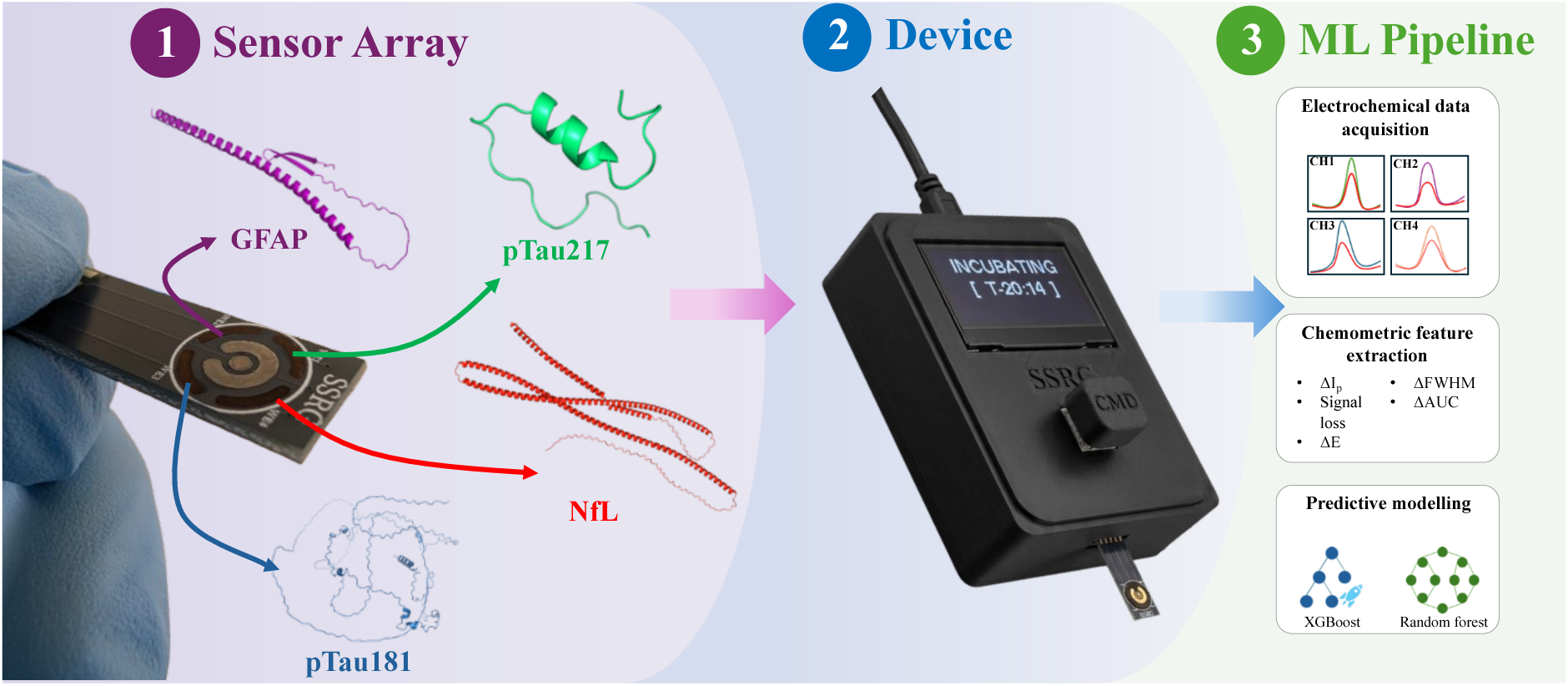
Schematic workflow and system architecture of the ML-enhanced multi-biomarker dementia diagnostic platform. Biosensor Array Interface: The four-channel PCB-based sensor array is held via a micro-patterned connector. The independent working channels are functionalized with tailored synthetic receptor matrices to target a comprehensive biomarker panel: WE1 for pTau217 (green), WE2 for GFAP (purple), WE3 for pTau181 (blue), and WE4 for NfL (red). Decentralized Hardware Integration: The sensor strip inserts directly into a custom, battery-powered handheld potentiostat featuring an integrated OLED display (showing real-time assay progress, e.g., “INCUBATING”) and local logging capabilities (scale bar: 10 mm). Acquisition & Calibration Pipeline: Electrochemical Data Acquisition: Real-time multi-channel Differential Pulse Voltammetry (DPV) sweeps are executed sequentially across the four channels to capture redox current suppression upon target binding. Chemometric Feature Extraction: High-dimensional descriptors are extracted from the baseline-corrected DPV curves—including Peak Height Difference (ΔI_p_), Signal Loss (%), Peak Potential Shift (ΔE), Delta Full Width at Half Maximum (ΔFWHM), and Delta Area Under the Curve (ΔAUC). Predictive Modelling: Cross-validated machine learning regressors (XGBoost and Random Forest) process the multi-dimensional feature space to resolve concentrations while mathematically suppressing crosstalk and biofouling interferences.

In addition, this multiplex format possesses significant operational advantages over single-biomarker assays in terms of overall cost, assay time, sample consumption, labor, and patient convenience. Rather than attempting a direct replacement or numerical conversion of established laboratory immunoassays, SMART-NeuroDx supports rapid point-of-care screening and triage using platform-specific thresholds.

To bypass the constraints of labour-intensive centralized systems like Single-Molecule Arrays (Simoa) or NuLISAseq, the SMART-NeuroDx platform integrates an array of Surface-Confined Redox-Active Molecularly Imprinted Polymers (Sc-RA-MIPs) onto highly reproducible printed circuit board (PCB) substrates to serve as robust, synthetic receptor analogues. This specific multi-channel SC-RA-MIP/AptaMIP/MIP matrix combination is reported here, delivering high sensitivity, selectivity across all target channels ^36^.

However, standard PCB surfaces (such as electroless nickel immersion gold (ENIG)) are highly vulnerable to pinhole degradation, mechanical wear, and severe non-specific biofouling in complex biological fluids like unconditioned human plasma, and high nickel reactivity in an electrochemical cell. To resolve these surface vulnerabilities, we implement a hardened electroplated gold barrier along with an interfacial modification utilizing highly porous gold (HPG) electrodeposited over the working electrode domain. The structural architecture of HPG provides a dual technical advantage, signal amplification via a massive electroactive surface area, and biofouling resilience ^37–40^. The interconnected microporous network of HPG introduces a definitive quasi-size-exclusion effect, physically obstructing larger macromolecular fouling species (such as lipids and bulky serum proteins) while allowing smaller target biomarkers to access the localized recognition cavities unobstructed^41^. Rather than altering the baseline material conductivity, this surface roughness acts as a geometric amplifier, maximizing the localized density of accessible redox sites and boosting analytical sensitivity.

For label-free and reagent-free signal transduction within this HPG-modified framework, we employ electropolymerized poly(phenol red) (pPhR) as the primary redox-active polymer backbone ^7^. Electropolymerization of the phenol red sodium salt yields a matrix with intrinsic, highly stable redox activity. Target binding directly modulates the internal charge-delocalization and electron transfer kinetics within the polymeric backbone, allowing binding events to be transduced cleanly as a direct suppression of the Faradaic current. This design eliminates the need for unstable external redox probes (such as ferri/ferrocyanide) or secondary reporting labels.

To fine-tune the mechanical stability and charge transport properties of these synthetic receptors, co-polymerization with functional monomers and crosslinkers is performed:

**Polypyrrole (pPy) :** Incorporated as a co-monomer to form a core composite matrix. The high electronic density and structural rigidity of the pyrrole ring introduce robust non-covalent *π*-*π* stacking and electrostatic coordination sites, securely locking target peptides into their biomimetic cavity geometries.

**Poly (ethylene glycol) (PEG):** Synthesized as a pyrrole-terminated PEG conjugate (PyPEG) crosslinker. The highly flexible, high-entropy polyether chains of the PEG framework extend into the liquid-electrode interface, coordinating water molecules to generate a physical hydration shell. This barrier suppresses non-specific protein adsorption and matrix fouling without blocking target diffusion into the embedded MIP cavities.

To target axonal injury rates with ultra-high affinity, the NfL channel utilizes a unique surface-confined aptamer-MIP hybrid (Sc-RA-AptaMIP/MIP) architecture ^36^. This hybrid design pairs a thiol-modified MN711 DNA aptamer with a robust polymer matrix ^11^. By enclosing the highly specific, nucleotide-level conformational recognition of the aptamer within the mechanical and thermal protective confinement of the synthetic polymer, the hybrid receptor achieves strong binding affinity and stability.

Despite the benefits of these localized matrices, fabricating a multi-analyte array on a single miniaturized substrate introduces a technical bottleneck: cross-contamination during the sequential functionalization of closely spaced channels. When a working electrode (WE) is immersed in a monomer-template precursor solution, adjacent channels are highly vulnerable to passive template fouling and non-specific polymer cross-deposition. We resolve this fundamental multi-channel fabrication barrier by introducing a potential-assisted sequential electropolymerization strategy that exploits the native isoelectric points (pI) of our comprehensive 4-biomarker panel. By systematically applying electrostatic-gated shielding potentials to adjacent unfunctionalized electrodes during each polymerization phase, we match the net surface charge of the co-existing template molecule at the operating pH. This electrostatically repels non-target templates, preventing passive cross-adsorption and preserving channel isolation across the array.

Furthermore, because complex biological matrices introduce overlapping chemical signals and minor baseline variations, chemometric machine learning (ML) integration is essential to make this platform a highly robust diagnostic utility ^42–47^. Raw differential pulse voltammetry (DPV) curves are converted into a high-dimensional functional feature space by extracting five primary descriptors: peak height difference (Δ*I_p_*), signal loss (%), peak potential shift (Δ*E*), delta area under the curve (ΔAUC), and delta full width at half maximum (ΔFWHM). These multi-dimensional descriptors are fed into robust regression models—such as Random Forest (RFR), Support Vector (SVR), and XGBoost Regressors—to mathematically isolate authentic target binding events from overlapping adjacent-channel signals or non-specific biofouling interferences ^42–47^.

Finally, to transition this ML-enhanced electrochemical array into a truly decentralized point-of-care system, the sensor is coupled with a custom-designed, battery-powered handheld potentiostat built around the STM32G491RCT6 microcontroller ^48^. This handheld unit integrates high-precision analogue peripherals, including high-speed 12-bit DACs for excitation waveform generation, low-noise operational amplifiers configured as transimpedance amplifiers (TIAs) with active auto-ranging circuitry, and synchronized 12-bit ADCs with real-time hardware oversampling to deliver benchtop-grade electrochemical performance in a pocket-sized, clinically deployment-ready format ^48^.

## Experimental

### Materials

Silver and gold brush plating solutions were procured from Spa Plating (Bath, UK). Gold (III) chloride, ammonium chloride, phenol red sodium salt (selected for its redox activity and compatibility with polymerization techniques), pyrrole, poly (ethylene glycol) diglycidyl ether (PEGDE), ferric chloride, potassium chloride, oxalic acid, and Tris(2-carboxyethyl) phosphine (TCEP) were sourced from Sigma-Aldrich (St. Louis, MO, USA). MN711 DNA aptamer was procured from Eurofins with HPCL purification and 3’ thiol modification^11^. Neurofilament light (NfL) was procured from CusaBio. Glial fibrillary acidic protein was procured from Abcam (Cambridge, UK). Printed circuit boards (PCBs) with 2 units (∼500 nm) electroless nickel immersion gold (ENIG) surface finish was obtained from JLC PCB (Shenzhen, China). Pooled human serum was procured from Sigma-Aldrich (St. Louis, MO, USA). Whole blood was collected from healthy donors by the University of Liverpool Biobank (LUB) in a K2EDTA tube and separated into plasma. The plasma was characterized for GFAP, pTau181, and NfL using SIMOA HD-X.

### Reagent preparation

100 mM and 10 mM Phosphate buffered saline and 100 mM HEPES were prepared pH buffered at 7.4 and 6.0 respectively. 10 mM Tris-HCl was prepared and pH buffered to 7.4 with KOH. 10 mM TCEP aliquots were prepared and frozen at −20 °C until further use. 20 mM AuCl_3_ and 2.5 M NH_4_Cl stocks were prepared for HPG deposition ^39,40^. 5 mM phenol red and 40 mM pyrrole stock solutions were prepared in 100 mM HEPES. Phenol red stock solution was degassed under a nitrogen stream for 15 minutes before use.

In a beaker 280 µL of DMSO was added along with 0.336 g of powdered KOH and stirred for 1 hour. 280 µL of pyrrole was added slowly under continuous stirring. After stirring for 1 hour 440 µL of PEGDE was added slowly under continuous stirring. The solution was allowed to stir overnight. The solution was then centrifuged at 20,000 RCF to separate the solids (KOH and pyrrole oligomers suspended in DMSO). The pale-yellow supernatant was separated and diluted with 100 mM HEPES (5 mM PEG equivalent) and stored at room temperature until further use.

The MN711 aptamer was primed by diluting the 2 µL of the 100 µM aptamer in 18 µL 10 mM PBS, followed by heating the sample at 95 °C for 10 minutes in a thermal cycler. The temperature was brought down to 25 °C at a gradient of 1.1%. 20 µL of 10 mM TCEP was added and the solution was incubated at 25 °C for 1 hour. 20 µL of this solution was added to 980 µL of Tris-HCl buffer and kept at 25 °C.

Monomer template precursors were separately for each biomarker. pTau 217 monomer template precursor was prepared by adding 200 µL of Phenol red 200 µL of PyPEG, 100 µL of 100 mM PBS, 475 µL of DI water and 25 µL of 0.2 mg mL^-1^ pTau217 peptide. GFAP monomer template precursor was prepared by 200 µL of Phenol red 200 µL of 100 mM HEPES, 100 µL of 100 mM PBS, 490 µL of DI water and 10 µL of 0.2 mg mL^-1^ GFAP. pTau181 monomer template precursor was prepared by 200 µL of Phenol red 200 µL of pyrrole, 100 µL of 100 mM PBS, 490 µL of DI water and 10 µL of 0.2 mg mL^-1^ pTau181 peptide. NfL monomer template precursor was prepared by 200 µL of Phenol red 200 µL of pyrrole, 100 µL of 100 mM PBS, 425 µL of DI water and 75 µL of 0.1 mg mL^-1^ NfL.

### Electrode preparation

The electrodes were designed with 4 identical islands as working electrodes (WE) and 1 common counter electrode (CE) and a common reference electrode (RE) in EasyEDA software. The electrodes were ordered as a panel of 20 x 10. The electrodes were cleaned in 70% ethanol by soaking for 15 minutes. The electrodes were then transferred to a water bath to remove any ethanol residue. Silver was electroplated on the CE and RE by performing 15 cycles of multistep potentiometry with a deposition cycle at −0.5 mA for 30 s and a stripping cycle at 0.1 mA. Gold electroplating was performed on the WEs by performing 25 cycles of multistep potentiometry with a deposition cycle at −1.0 mA for 30 s and a stripping cycle at 0.2 mA. CE and RE were treated with 20 µL of 3 M KCl for 30 seconds and the RE was modified to Ag/AgCl by dropping 2 µL of 100 mM FeCl_3_. 20 mM AuCl_3_ and 2.5 M NH_4_Cl were mixed in a 1:1 ratio and soft gold deposition was performed using 10 cycles cyclic voltammetry from 0.8 to 0.3 V (Vs Ag/AgCl) at 0.1 V/s. Electroporation was performed by running chronoamperometry at −1.2 V (Vs Ag/AgCl) for 60 seconds. The electrodes were then rinsed and dried.

### Sequential functionalization

Before functionalization the electrodes were electrochemically cleaned in 10 mM H_2_SO_4_ by performing 5 cycles of cyclic voltammetry from −0.4 to 0.6 V (Vs Ag/AgCl) ^6^.

10 µL of the aptamer solution was added to the WE4 and incubated on a wet paper towel ^36^. 10 µL of 6.1 µM of NfL was added to the WE4 to bind to the immobilized aptamer ^36^. The pI of each template molecule was computed using swissprot and are listed in table 1 ^49,50^.

**Table 1:** Table showing the isoelectric point (pI) and gating potential during polymerization for each biomarker.

| Template molecule | pI | Gating potential |
| --- | --- | --- |
| NfL | 5.83 | -0.3 V (Vs Ag/AgCl) |
| pTau181 | 10.80 | 0.2 V (Vs Ag/AgCl) |
| GFAP | 4.60 | -0.3 V (Vs Ag/AgCl) |
| pTau217 | 11.00 | 0.2 V (Vs Ag/AgCl) |

100 µL of the monomer template precursor was added to the electrode and sequential functionalization was performed using 18 cycles (20 for GFAP) of cyclic voltammetry from 0.3 to 0.8 V (Vs Ag/AgCl) at 0.1 V/s. Template removal was performed by incubating the electrodes in 50 mM oxalic acid at 4 °C overnight followed by a 15-minute soak in 10 mM PBS.

## Electrochemical Characterization and Analytical Performance Validation

All electrochemical metrics were evaluated using a CHI1030a potentiostat configuration via differential pulse voltammetry (DPV) and cyclic voltammetry (CV). DPV scans were swept from −0.2 V to 0.5 V with an amplitude of 50 mV, 20 mV step potential, pulse width of 100 ms, and pulse period of 500 ms to evaluate the oxidation current profile of the redox-active polymer backbones.

### Dose-Response Evaluation

To ground the calibration in authentic sample composition, the endogenous plasma concentrations of GFAP, pTau181, and NfL in the pooled healthy-donor plasma were first determined on a Simoa HD-X analyzer. These reference baselines defined the spiking scheme, so that each calibration curve spanned the physiologically relevant range above the measured native background; reported plasma concentrations for these three markers therefore represent the Simoa-determined baseline plus the added spike. pTau217 was spiked across its calibration range in each matrix; its endogenous plasma concentration was not separately quantified by the reference method. Calibration profiles for each working electrode (WE1–WE4) were recorded in triplicate by spiking known concentrations of the respective biomarkers into 10 mM PBS, unprocessed human plasma, and serum.

Individual calibration profiles for each working electrode (WE1–WE4) were recorded in triplicate by spiking varying concentrations of the respective biomarkers into 10 mM PBS, unprocessed human plasma and serum.

### Selectivity factor

The selectivity of the Sc-RA-MIP was evaluated for each channel using the slope of dose-response in PBS against plasma and serum using equation 1:

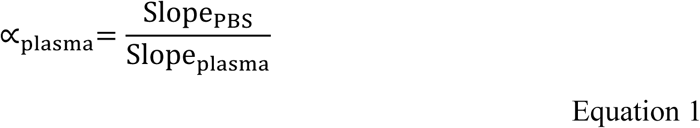

### LOD/LOQ Determination

The limit of detection (LOD) and limit of quantification (LOQ) for each biomarker channel were calculated using the standard deviation of the blank signal (*σ*, *n* = 18) and the slope of the linear range (*S*) using equations 2 and 3:

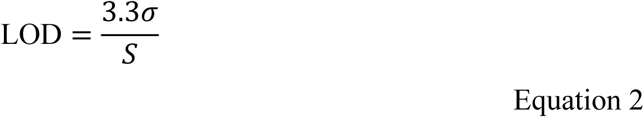

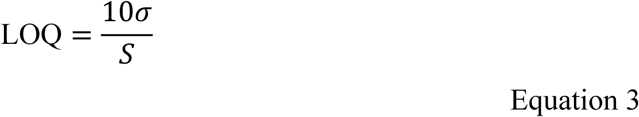

RSD calculations: The relative standard deviation (RSD) was calculated using the standard deviation (*σ*, *n* = 18) of the population and µ is the mean of the population at half maxima, using equation 4:

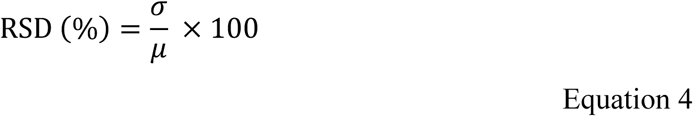

## Cross-Reactivity and Interference

### Cross-Reactivity Testing

To verify individual channel separation and ensure negligible sensor-to-sensor communication, cross-reactivity studies were performed in 10 mM PBS (pH 7.4). Each individual target pTau217 (3 pg ⋅ mL^−1^), GFAP (300 pg ⋅ mL^−1^), pTau181 (15 pg ⋅ mL^−1^), and NfL (20 pg ⋅ mL^−1^) was spiked into the testing cell separately. The Faradaic responses were tracked concurrently across all four working electrode channels (WE1-WE4) using sequential Differential Pulse Voltammetry (DPV) to confirm the absence of electrochemical or physical crosstalk between adjacent sensing microstructures.

### Interference Assays

To evaluate the platform’s clinical durability and selectivity under challenging physiological backgrounds, the multi-channel arrays were evaluated against complex biological background matrices. Interference profiles were established by systematically testing each channel’s target signal response in solutions containing a massive, pathological background of Bovine Serum Albumin (BSA, 35 mg ⋅ mL^−1^), alongside relevant co-existing systemic inflammatory and neurodegenerative targets. These included

Brain Natriuretic Peptide (BNP, 200 pg ⋅ mL^−1^), Interleukin-6 (IL-6, 200 pg ⋅ mL^−1^), Interleukin-18 (IL-18, 400 pg ⋅ mL^−1^), pathological C-Reactive Protein (pCRP, 30 µg ⋅ mL^−1^), and Amyloid-beta 1-40 (A*β*_1-40_, 400 pg ⋅ mL^−1^) ^51–57^.

The sensor array was incubated in these mixed analyte-interferent matrices at 20°C for 30 minutes, followed by a brief, gentle washing step with ultra-pure water and PBS to remove weakly bound, non-specific proteins before DPV signal acquisition. The percentage of signal loss was calculated and compared directly to baseline measurements in PBS to verify high-fidelity signal collection under competitive binding environments.

## Chemometric Feature Extraction and Machine Learning Pipeline

To maximize the predictive accuracy of the multi-channel array and systematically suppress the confounding effects of electrochemical crosstalk and complex matrix interference, a targeted chemometric feature extraction pipeline was established prior to model training. Raw differential pulse voltammetry (DPV) curves from each working electrode (WE1-WE4) were converted into a high-dimensional functional feature space. Instead of relying solely on absolute peak current measurements, five primary chemometric descriptors were mathematically extracted from the baseline-corrected voltammetric profiles:

### Peak Height Difference (Δ*I*_p_, *μ*A)

The absolute change in the Faradaic peak current relative to the clean baseline buffer response, capturing the primary scale of cavity occupancy.

**Signal Loss (**%)

The normalized percentage decrease in the Faradaic current, calculated as:

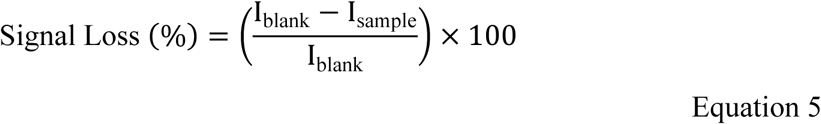

providing a matrix-invariant scaling metric across different sensor chips.

### Peak Shift (Δ*E*_p_, mV)

The lateral displacement of the peak potential, tracking alterations in electron transfer kinetics and internal charge-delocalization caused by template binding or microenvironmental matrix variations.

### Delta Area Under the Curve (ΔAUC)

The differential integrated area beneath the voltammetric peak profile, which captures aggregate charge transfer modifications and compensates for asymmetric peak distortions.

### Delta Full Width at Half Maximum (ΔFWHM)

The change in the peak width at half-maximum amplitude, acting as a sensitive indicator of underlying changes in the apparent electron transfer coefficient (*α*) and distinguishing target-specific binding from non-specific macromolecular surface adsorption.

By capturing these multi-dimensional peak variations simultaneously, the extracted feature set allowed the downstream algorithms to mathematically isolate authentic template binding events from overlapping adjacent-channel signals or non-specific biofouling interferences. Following chemometric processing, independent regression models were assigned to each distinct biomarker channel. Model selection was systematically performed by evaluating and benchmarking three robust regression architectures: Random Forest Regressor (RFR), Support Vector Regression (SVR) with a radial basis function (RBF) kernel, and Extreme Gradient Boosting (XGBoost) Regressor. For each biomarker model, hyperparameter tuning and optimization were executed via an automated grid search cross-validation (GridSearchCV) strategy. The search space evaluated key structural hyperparameters, including:

#### RFR

Number of estimators, maximum tree depth, and minimum sample split thresholds.

#### SVR

Penalty parameter (*C*), epsilon (*ε*), and kernel coefficient (*γ*).

#### XGBoost

Learning rate (*η*), maximum depth, subsample ratio, and regularization weights (*α* and *λ*).

The total number of collected voltammetric data points varied across the four biomarker channels (WE1-WE4) due to the progressive development and timeline of the multi-channel database over a two-year experimental period. Channels targeting pTau181 (*n* = 11,320), NfL (*n* = 9,260), and GFAP (*n* = 7,862) were integrated during initial platform development (currently under peer review), while the pTau217 channel (*n* = 980) represents the latest addition to the diagnostic panel. Despite dataset size variations, all four channel models were trained using identical 5-fold cross-validation and hyperparameter optimization workflows to ensure balanced, overfit-resistant generalization.

To prevent overfitting and ensure robust model generalization across independent sensor chips, the dataset for each channel was partitioned using a standard train-test split, followed by a rigorous five-fold cross-validation scheme applied exclusively to the training subset. The predictive performance of the candidate algorithms was assessed using root mean square error (RMSE) and coefficient of determination (*R*^2^) validation metrics. The optimized model exhibiting the lowest cross-validated RMSE was subsequently frozen and validated against the holdout test set to deliver high-fidelity concentration predictions directly from raw electrochemical data profiles.

#### Safety

Pyrrole is harmful and is a skin and respiratory irritant; it was handled neat and in dimethyl sulfoxide (DMSO) under local exhaust ventilation. The PyPEG synthesis, which combines pyrrole, powdered potassium hydroxide, and DMSO, was carried out with particular care, as DMSO promotes dermal absorption of co-dissolved species and potassium hydroxide is corrosive. Gold (III) chloride, ferric chloride, oxalic acid, and dilute sulfuric acid are corrosive or harmful and were handled with nitrile gloves, eye protection, and appropriate containment; PEGDE is a skin sensitizer. Phenol red stock was degassed under nitrogen in a ventilated area to avoid nitrogen accumulation in enclosed spaces.

Electrodeposition, soft-gold deposition, and electroporation steps involve applied currents and potentials and were performed on isolated, current-limited instrumentation. Human whole blood, plasma, and pooled serum were treated as potentially infectious and handled under biosafety level 2 practice, with waste inactivated and disposed of in accordance with institutional biosafety and COSHH procedures. No unexpected hazards were encountered.

## Results and Discussion

### Analytical Validation and Dose-Response Profiles

To evaluate the clinical utility of the integrated PCB sensor array, the analytical performance of each Sc-RA-MIP-modified working channel (WE1-WE4) was systematically validated across physiologically relevant clinical concentrations. Quantitative analysis was performed by tracking the percentage signal loss of the intrinsic Faradaic current via differential pulse voltammetry (DPV) as a function of target concentration (representative DPVs shown in **Figure S2**). Global dose-response saturation profiles were fitted to the Hill model:

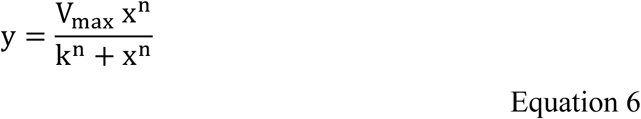

where y represents the percentage signal loss, *x* is the analyte concentration, *V*_max_ is the maximum achievable signal suppression at saturation, k is the apparent dissociation constant, and *n* is the Hill coefficient.

To establish the limits of detection (LOD) and quantification (LOQ), linear regression was applied strictly to the low-concentration, linear regime of each biomarker’s dataset. While global data adhere to the non-linear Hill model, the initial analytical sensitivity (*S*) was extracted directly from these localized linear ranges. This dual-modeling approach ensures statistically rigorous trace-level detection thresholds while capturing global saturation dynamics at higher clinical concentrations.

These calibrations were performed directly in unprocessed plasma and serum rather than in buffer alone, so they represent matrix-based recovery in the intended clinical sample type. For GFAP, pTau181, and NfL, the spiking range was anchored to endogenous concentrations independently determined by Simoa HD-X, linking the calibration to a reference-quantified baseline. Across the four channels, dose-response slopes in plasma and serum remained close to those in PBS, with selectivity factors between 0.84 and 1.28 (Table 2), indicating that the reagent-free interface recovers spiked analyte with only modest, quantifiable matrix attenuation. These results establish fit-for-purpose analytical performance in authentic biofluids.

**Table 2:** Analytical Performance Metrics of the Multiplexed Biosensor Array.

| Biomarker | WE | LOD<br>(pg mL<sup>-1</sup>) | LOQ<br>(pg mL<sup>-1</sup>) | Range<br>(pg mL<sup>-1</sup>) | RSD<br>(%) | $\alpha_{\text{plasma}}$ | $\alpha_{\text{serum}}$ |
| --- | --- | --- | --- | --- | --- | --- | --- |
| pTau217 | WE1 | 0.087 | 0.26 | 0-10 | 5.07 | 0.87 | 0.85 |
| **GFAP** | WE2 | 31.3 | 94.84 | 0-600 | 3.23 | 1.14 | 1.18 |
| **pTau181** | WE3 | 0.98 | 2.96 | 0-30 | 5.34 | 1.26 | 1.28 |
| **NfL** | WE4 | 5.83 | 17.66 | 0-100 | 3.95 | 0.91 | 0.84 |

*Keys: WE: Working electrode channel; LOD: Analytical limit of detection; LOQ: Analytical limit of quantification; Range: Demonstrated dynamic range in K2EDTA Plasma; RSD: Relative standard deviation; $\alpha$ : Selectivity factor calculated using Equation 1.*

**Table 3:**
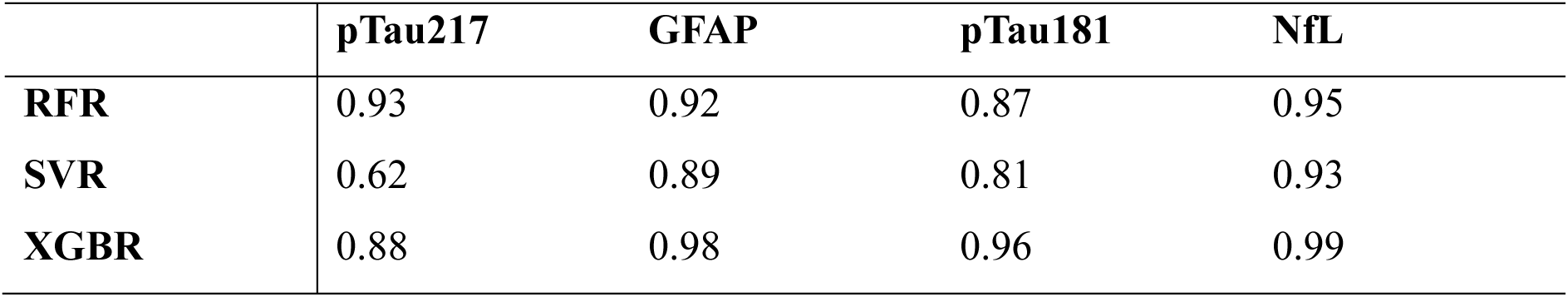
Summary of model performance for each biomarker.

### Channel-Specific Profiles

#### pTau217 (WE1)

Demonstrating sub-picogram sensitivity, WE1 achieved an LOD of 0.087 pg ⋅ mL^−1^ and LOQ of 0.264 pg ⋅ mL^−1^ across a 0-10 pg ⋅ mL^−1^ range (Adj. *R*^2^ = 0.99). The Hill coefficient (*n* = 1.59) in plasma indicates positive competitive binding profile, where PEG chains prevent non-specific fouling while leaving high affinity nanocavities accessible. The channel maintained stable reproducibility (RSD = 5.07%) and a matrix selectivity factor *α_plasma_* of 0.87 and *α_serum_* of 0.85 (**Figure 2A**).

**Figure 2.**
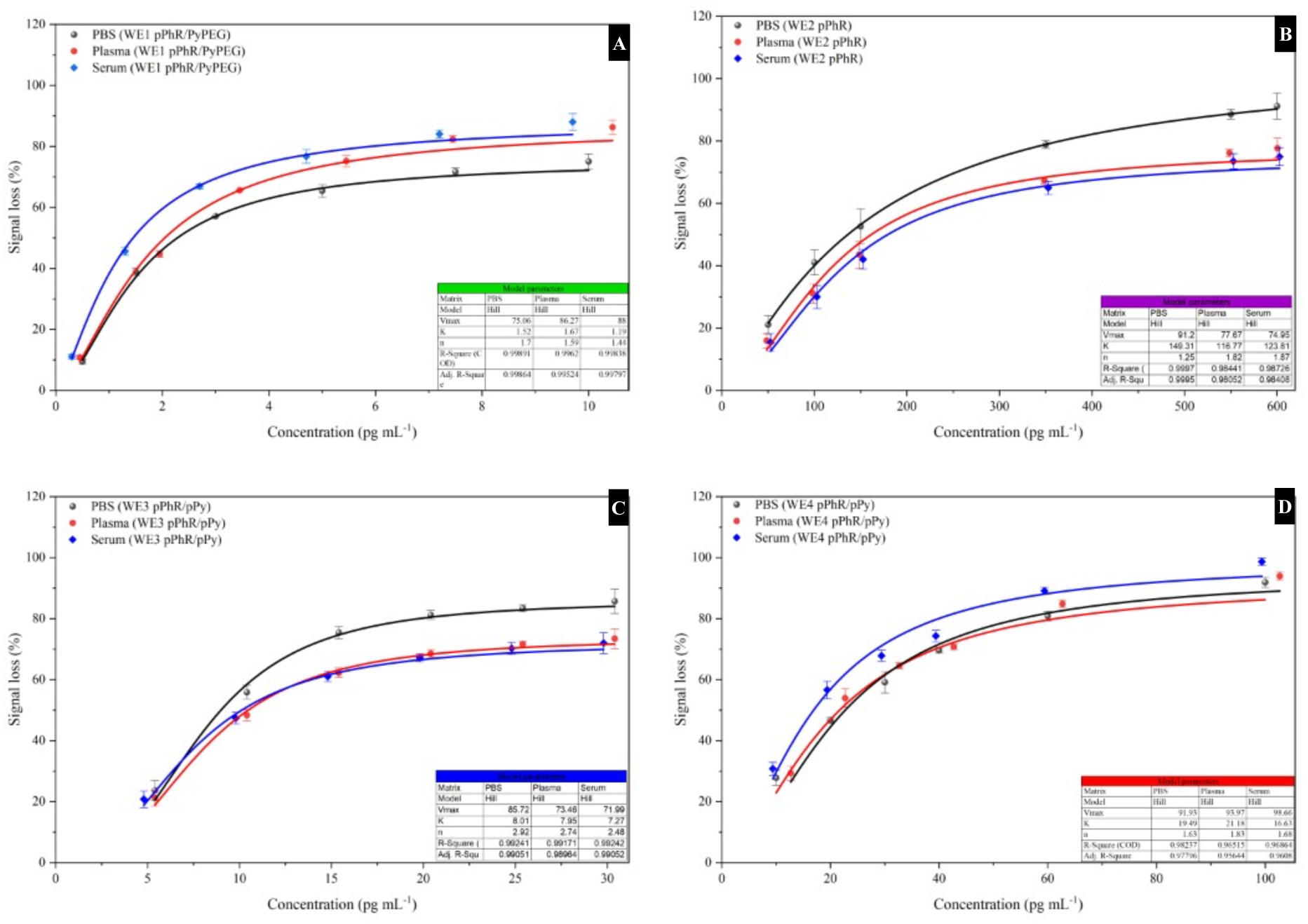
**Multi-channel non-linear calibration curves and Hill-model fittings for the multiplexed biosensor array in buffer and complex clinical matrices. Dose-response profiles comparing Faradaic signal loss (%) in 10 mM PBS (black spheres), raw human plasma (red circles), and raw human serum (blue diamonds) are presented for:** **(A) pTau217** (WE1) spanning a range of 0–10 pg mL⁻¹; **(B) GFAP** (WE2) spanning a range of 0–600 pg mL⁻¹; **(C) pTau181** (WE3) spanning a range of 0–30 pg mL⁻¹; and **(D) NfL** (WE4) spanning a range of 0–100 pg mL⁻¹.

#### GFAP (WE2)

Monitoring astrocytic neuroinflammation, WE2 exhibited close fit to the Hill model (Adj. *R*^2^ = 0.999) over a broad 0-600 pg ⋅ mL^−1^ range (LOD = 31.3 pg ⋅ mL^−1^, LOQ = 94.839 pg ⋅ mL^−1^). The calculated Hill coefficient (*n* = 1.82) points to strong cooperative binding dynamics within the homogeneous pPhR Sc-RA-MIP matrix, accompanied by a stable complex matrix response (α_plasma_ = 1.14, α_serum_ = 1.18) and an RSD of 3.23% (**Figure 2B**).

#### pTau181 (WE3)

Tracking classic tauopathy hallmarks across 0-30 pg ⋅ mL^−1^ (Adj. *R*^2^ = 0.99), WE3 reached an LOD of 0.98 pg ⋅ mL^−1^ and LOQ of 2.969 pg ⋅ mL^−1^. The pPhR/pPy matrix yielded a Hill coefficient (*n* = 2.74), reflecting extremely cooperative site occupancy within the cavities (RSD = 5.34%, α_plasma_ = 1.26, α_serum_ = 1.28). The high Hill coefficient suggests that the polymer matrix consists of at least 3 to 4 interacting binding sites grouped closely together in the local microenvironment ^58^ (**Figure 2C**).

#### NfL (WE4)

Leveraging a surface-confined aptamer-MIP hybrid (Sc-RA-AptaMIP/MIP), WE4 yielded an LOD of 5.83 pg ⋅ mL^−1^ and LOQ of 17.665 pg ⋅ mL^−1^ across 0-100 pg ⋅ mL^−1^ (Adj. *R*^2^ = 0.977). The hybrid MN711 aptamer and pPhR/pPy matrix produced positive cooperativity (*n* = 1.31), low matrix vulnerability (α_plasma_ = 0.91, α_serum_ = 0.84), and high operational precision (RSD = 3.95%) (**Figure 2D**).

Dose-response was performed on 24 individual devices (4 replicates) for each sample matrix. Error bars represent standard error.

Furthermore, the long-term shelf-life of pPhR-based Sc-RA-MIP matrices over a 1-year storage period has been previously reported, demonstrating high operational resilience without strict cold-chain dependence ^59^.

### Selectivity and Cross-Reactivity Evaluation in Complex Matrices

Interference and cross-reactivity challenges were systematically conducted in standardized 10 mM PBS rather than native human plasma or serum to avoid confounding baseline signals from endogenously circulating target biomarkers and inflammatory proteins, which would otherwise require extensive, separate immunoassay quantification. To evaluate the analytical specificity and potential cross-reactivity of the multiplexed diagnostic array, interference studies were systematically executed. Each optimized working electrode channel (WE1-WE4) was exposed to its respective target biomarker in the presence of a wide range of highly abundant blood-borne proteins, inflammatory markers, and competing neurodegenerative targets. The target and non-target concentrations were chosen to carefully match or exceed high physiological and pathological clinical thresholds:

**Target Biomarkers:** pTau217 (3 pg ⋅ mL^−1^), pTau181 (15 pg ⋅ mL^−1^), GFAP (300 pg ⋅ mL^−1^), and NfL (20 pg ⋅ mL^−1^).

**Endogenous Neuro- & Cardiovascular Interferents:** Amyloid-beta 1-40 (A*β*_1-40_, 400 pg ⋅ mL^−1^) and Brain Natriuretic Peptide (BNP, 200 pg ⋅ mL^−1^).

**Systemic Inflammatory Mediators:** Interleukin-6 (IL-6, 200 pg ⋅ mL^−1^), Interleukin-18 (IL-18, 400 pg ⋅ mL^−1^), and pathological C-Reactive Protein (pCRP, 30 *μ*g ⋅ mL^−1^).

**Abundant Plasma Matrix Background:** Bovine Serum Albumin (BSA, 35 mg ⋅ mL^−1^).

The electrochemical response for each channel was quantified as a percentage of signal loss (%, representing the current suppression caused by target capture within the respective molecularly imprinted polymer cavities) and compared directly to a baseline blank control (PBS), as displayed in **Figure 3A–D**.

**Figure 3.**
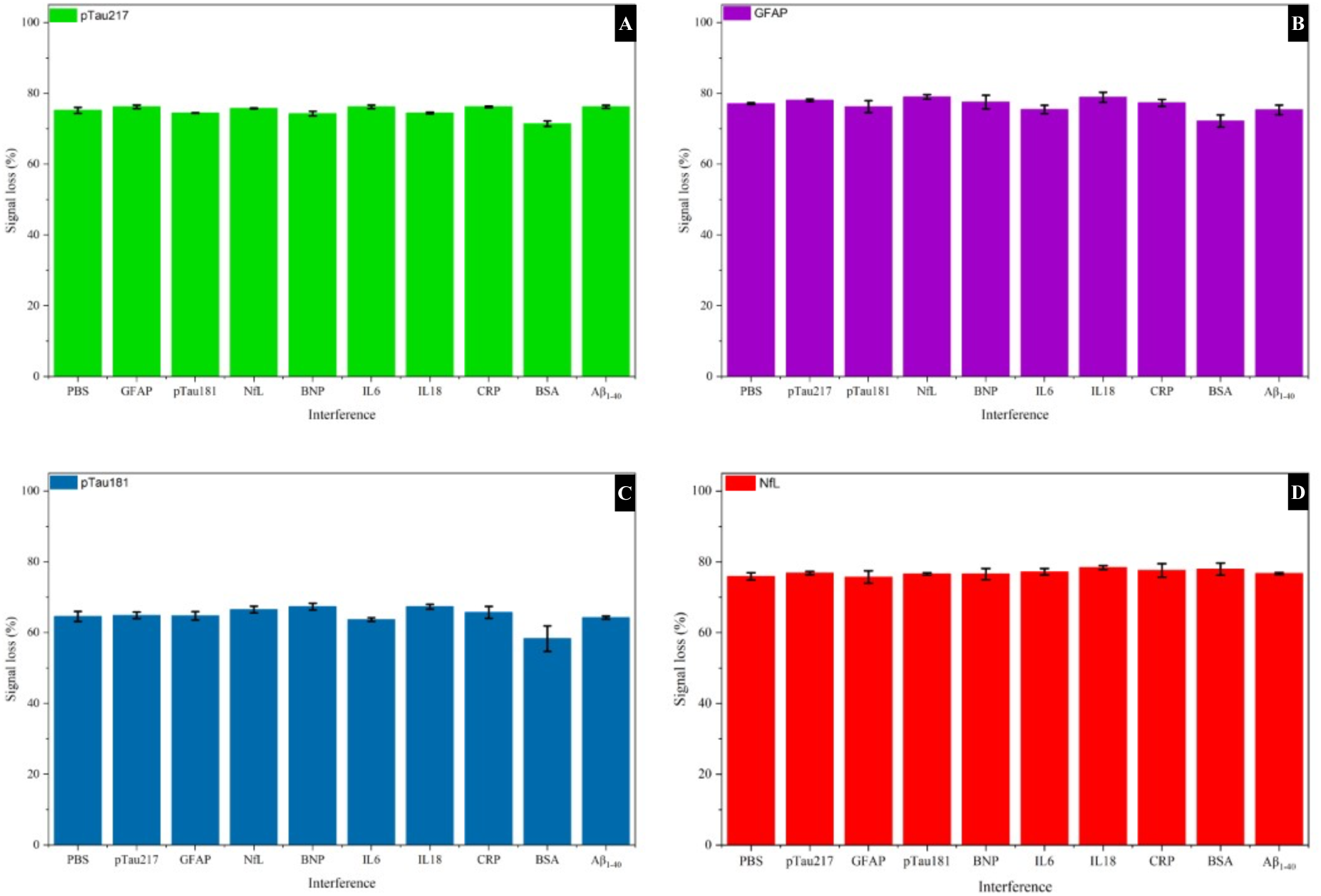
Selectivity and cross-reactivity profiles of the multi-channel biosensor array under complex physiological and competitive background matrices. The percentage of Faradaic signal loss (%) is plotted for each functionalized working electrode channel in the presence of its respective target biomarker alongside various high-concentration clinical interferents: **(A) pTau217** (WE1, green) evaluated with its target at 3 pg ⋅ mL^−1^;χσβαρλινε **(B) GFAP** (WE2, purple) evaluated with its target at 300 pg ⋅ mL^−1^; **(C) pTau181** (WE3, blue) evaluated with its target at 15 pg ⋅ mL^−1^; and **(D) NfL** (WE4, red) evaluated with its target at 20 pg ⋅ mL^−1^.

The pTau217 channel demonstrated high selectivity, maintaining a highly stable signal loss profile of approximately 74.5% to 76.5% across nearly all co-existing interferents, compared to the 75.1% baseline response in PBS (**Figure 3A**). The maximum deviation was recorded in the presence of an extreme overload of BSA (35 mg ⋅ mL^−1^), which caused a minor decrease in signal loss to 71.6% (a negligible recovery deviation of under 4.7%). This demonstrates that the protective, hydrated PyPEG polyether chains on WE1 successfully suppress non-specific biofouling even under highly concentrated protein conditions.

The GFAP channel showed strong resilience to co-existing proteins, with signal loss values consistently hovering around 75.5% to 79.0% against the PBS control (77.1%) (**Figure 3B**). No significant cross-reactivity was observed with the closely related structural protein fragments or tau isoforms. The lowest signal loss value was observed under the massive BSA background (72.3%, representing a minor −6.2% deviation), validating the structural stability of the pure pPhR cavity configuration.

The pTau181 sensing channel exhibited stable electrochemical behaviour, with signal loss values closely clustered around the PBS baseline of 64.7% (**Figure 3C**). Cross-analyte testing against the other major tauopathy target (pTau217) yielded 65.2% signal loss, demonstrating clear discrimination between the two hyperphosphorylated tau variants at different residues. Under the high-background BSA interference, the signal loss shifted slightly to 58.3%, remaining well within acceptable analytical tolerances (relative standard deviation < 9.8%) for blood-based clinical diagnostic assays.

The NfL sensing channel achieved the most consistent response across all tested matrices, with signal loss values tightly constrained within a narrow range of 75.2% to 78.5% relative to the 75.9% PBS control (**Figure 3D**). Crucially, the channel showed virtually no deviation when exposed to the massive BSA background (78.0%) or highly elevated levels of acute-phase proteins like CRP (77.5%). This performance is attributed to the rigid, highly customized binding cavities within the electropolymerized pPhR/pPy composite matrix, which selectively recognize the structural folds of neurofilament light chain while rejecting highly abundant blood-derived interferents.

Overall, the maximum signal deviations across all four optimized microelectrodes remained well below the critical ±10% threshold under both high-concentration biomarker cross-contamination and extreme protein background conditions (specifically, 35 mg ⋅ mL^−1^ BSA). These results mathematically confirm the selectivity, anti-biofouling performance, and overall diagnostic reliability of this multi-channel point-of-care platform for clinical blood plasma screening.

Interference and cross-reactivity study was performed in 10 mM PBS on 36 individual devices (4 replicates). Error bars represent the standard error.

### Surface characterization

Raman chemical mapping and FTIR-ATR spectroscopy confirmed the expected functional-group chemistry of each polymer matrix and the strict spatial confinement of each formulation to its designated channel. FTIR-ATR spectra, are provided in the Supporting Information (**Figures S4**).

### Raman Spectroscopic Characterization of the Conjugated Polymer Matrices

Raman spectroscopy (400–2000 cm⁻¹) was performed on WE1 (pPhR/PyPEG, 1:2 pPhR: pyrrole via bis-pyrrole PEG crosslinker), WE2 (pPhR only), and WE3/WE4 (pPhR/pPy, 1:8, replicates).

#### Spatial confinement (**Figure 4A–B**)

The Raman chemical map shows clean, boundary-matched chemical assignment for each electrode (WE1 white, WE2 red, WE3/WE4 blue) against the gold crescent geometry, confirming film growth is confined to the targeted electrode area with no cross-channel contamination.

**Figure 4.**
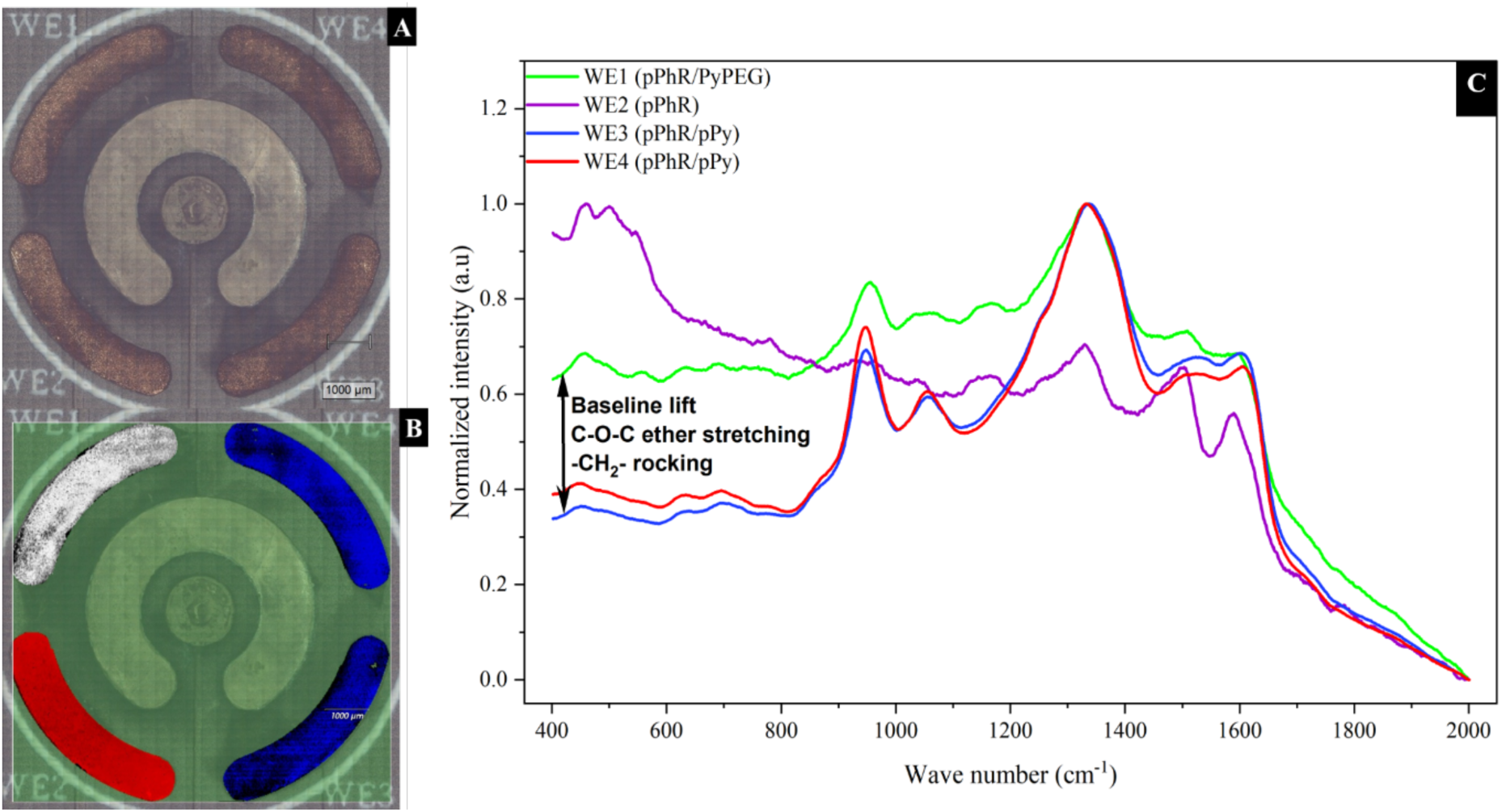
Spatial Raman chemical mapping and channel-specific spectral profiles of the four-channel sensor array. **(A)** Optical micrograph showing the baseline morphology of the four-channel sensor array platform (scale bar: 1000 μm). **(B)** Confocal Raman chemical map illustrating the strict spatial confinement and uniform distribution of the electrodeposited polymer matrices across the independent working channels: **NOTE: Figure S4B uses independent colour scheme due to fixed software output: WE1** (white), **WE2** (red), **WE3** (blue), and **WE4** (blue). **(C)** Normalized Raman point-spectra extracted from each individual channel over the 400 to 2000 cm^−1^ wavenumber region. The arrow notes the key spectral baseline lift, C-O-C ether stretching, and −CH_2_ − rocking modes unique to the pegylated PyPEG crosslinked matrix on **WE1**.

#### WE2 (pPhR)

Shows only pPhR aromatic backbone vibrations; the pyrrole-diagnostic 945 and 1340 cm⁻¹ bands are absent (heights 0.008 and 0.036, noise-level), confirming a clean pyrrole-free reference (**Figure 4C**).

#### WE3/WE4 (1:8 pPhR: pPy)

The pyrrole fingerprint dominates, with close replicate agreement (945 cm⁻¹: 0.213/0.246; 1340 cm⁻¹: 0.287/0.306; within 7–13%). Key bands: 945/1050 cm⁻¹ (pyrrole ring deformation/C–H bending), 1340 cm⁻¹ (dominant skeletal C–C stretch), 1500–1600 cm⁻¹ (C=C backbone stretch of doped pyrrole) (**Figure 4C**).

#### WE1 (1:2 pPhR: pPy, PyPEG)

Despite a quarter of the pyrrole content of WE3/WE4, WE1 shows clearly resolved 945 and 1340 cm⁻¹ bands (0.108, 0.134), confirming both pyrrole termini of PyPEG are electroactive and conjugated into the backbone. Normalized per pyrrole equivalent, WE1’s 945 cm⁻¹ intensity (∼0.054) is roughly double that of WE3/WE4 (∼0.027– 0.031) indicating PyPEG incorporates pyrrole more efficiently per unit than free-pyrrole excess, consistent with it acting as a defined crosslinking node rather than a simple additive. The elevated, broadened baseline beneath WE1’s pyrrole bands reflects PEG C–O–C ether/– CH₂– background rather than reduced pyrrole incorporation (**Figure 4C**).

### Hardware Instrumentation and Benchtop Validation

The four-channel assay was implemented on a custom, battery-powered STM32G491-based handheld potentiostat providing excitation, per-channel transimpedance amplification with active auto-ranging, synchronized acquisition and on-device data logging. Baseline differential pulse voltammograms recorded on the handheld unit matched those from a commercial CHI1030A workstation across all four channels, confirming benchtop-equivalent fidelity (**Figure S3**). The full circuit architecture (block diagram) is detailed in the Supporting Information (**Figure S5**).

### Model Selection and Machine Learning Validation

To mathematically isolate authentic target biomarker binding events from non-specific matrix interferences and overlapping cross-channel signals, a comprehensive chemometric machine learning (ML) framework was implemented. Five multi-dimensional functional descriptors derived from raw differential pulse voltammetry (DPV) curves were used to train three robust regression algorithms: Random Forest Regressor (RFR), Support Vector Regressor (SVR), and XGBoost Regressor (XGBR). Evaluation of the baseline *R*^2^ scores across the multi-channel panels calculated following hyperparameter optimization using GridSearchCV with 5-fold cross-validation directed the final model selection layout to guarantee optimal prediction accuracy for each independent diagnostic channel:

Following model selection, 100 sample validation sets were extracted to construct actual versus predicted concentration correlation plots, mapping the direct translation of high-dimensional voltammetric features into absolute biomarker metrics.

#### **A.** pTau217 Verification (WE1 — RFR Architecture)

For the trace-level pTau217 channel, the Random Forest Regressor (RFR) was chosen as the primary deployment architecture based on its superior training performance (*R*^2^ = 0.93).

Evaluated over a validation subset drawn from a comprehensive population of 980 total data points, the RFR algorithm achieved tight concentration mapping across the narrow 0 to 10 pg ⋅ mL^−1^ dynamic range (**Figure 5A**). Linear regression analysis of the actual vs. predicted distribution yielded a high coefficient of determination (*R*^2^ = 0.94137) paired with an adjusted *R*^2^ of 0.94077. The optimized slope (1.00837 ± 0.02542) and negligible intercept (−0.06557 ± 0.14714 pg ⋅ mL^−1^) confirm high prediction fidelity and minimal systemic skew within the ternary pPhR/PyPEG protective matrix.

**Figure 5.**
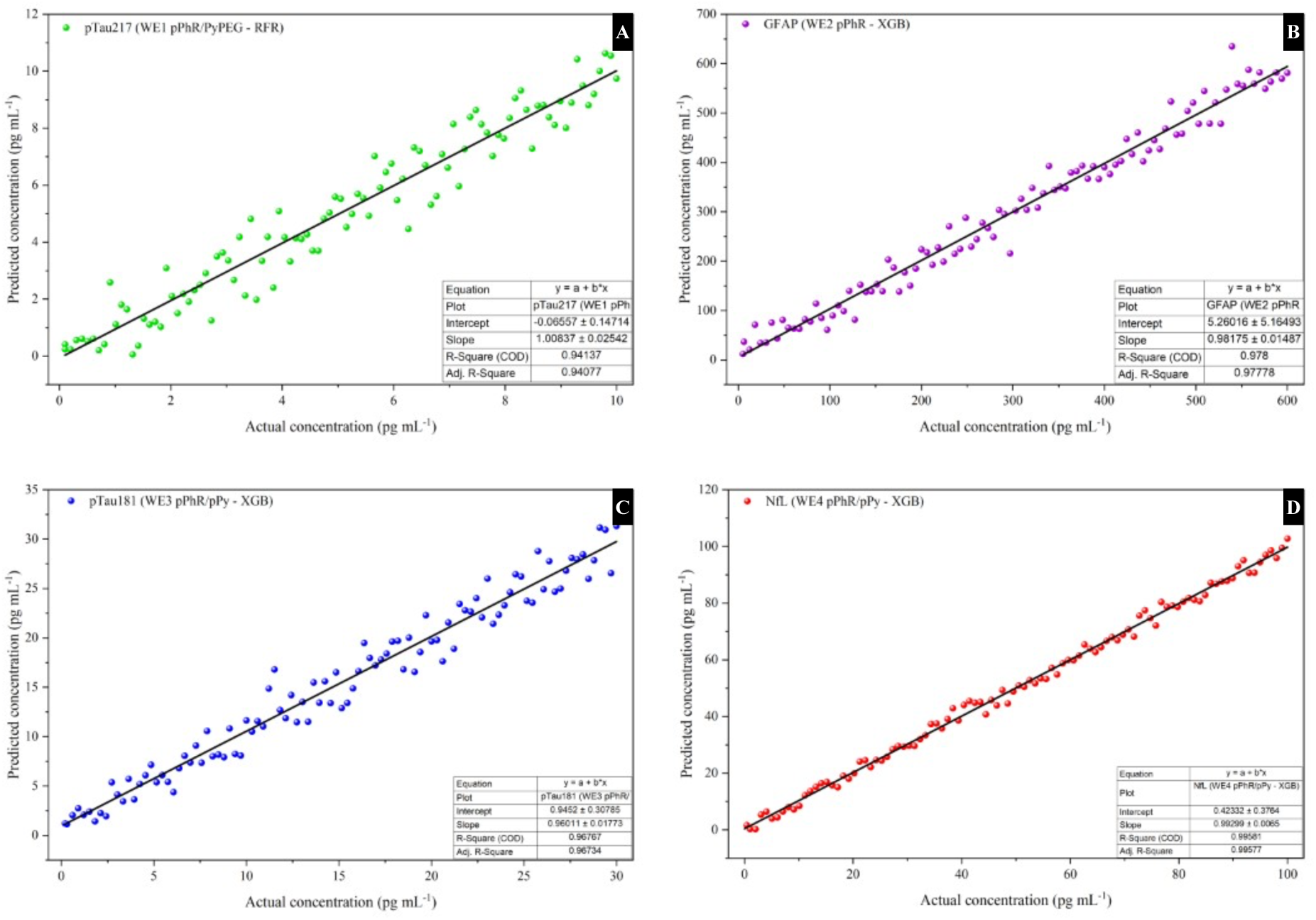
Actual versus chemometric model predicted concentration correlation plots across the four independent biosensor channels. Linear correlation profiles mapping actual concentrations against machine learning predicted concentration values derived from high-dimensional functional DPV feature extraction. Plots represent 100 validation samples evaluated per channel after screening optimal regression architectures: **(A) pTau217** (WE1) processed via Random Forest Regression (RFR) across a 0 to 10 pg ⋅ mL^−1^ range; **(B) GFAP** (WE2) processed via XGBoost Regression (XGB) across a 0 to 600 pg ⋅ mL^−1^ range; **(C) pTau181** (WE3) processed via XGBoost Regression (XGB) across a 0 to 30 pg ⋅ mL^−1^ range; and **(D) NfL** (WE4) processed via XGBoost Regression (XGB) across a 0 to 50 pg ⋅ mL^−1^ range.

#### **B.** GFAP Verification (WE2 — XGBoost Architecture)

The expanded dynamic range required for tracking neuroinflammatory GFAP (0 to 600 pg ⋅ mL^−1^) was mapped using the XGBoost Regressors (XGBR), which outperformed competing models with a selection *R*^2^ score of 0.98. Extracted from a broad pool of 7,862 data points, the 100-sample validation plot displays superb linearity across the entire analytical range (**Figure 5B**). The fitted regression curve generated an *R*^2^ value of 0.978 (Adj. *R*^2^ = 0.97778). The resolved slope (0.98175 ± 0.01487) tightly tracks the ideal unity line, accompanied by a small software calibration offset intercept of 5.26016 ± 5.16493 pg ⋅ mL^−1^, proving that the tree-boosting algorithm effectively handles the minor matrix attenuation typical of the pure pPhR homopolymer interface.

#### **C.** pTau181 Verification (WE3 — XGBoost Architecture)

The pTau181 classic tauopathy index channel achieved an optimal model selection *R*^2^ score of 0.96 utilizing the XGBR framework. Validated using 100 targeted testing arrays chosen from a substantial dataset of 11,320 operational points, the model successfully deconvoluted complex signals across a 0 to 30 pg ⋅ mL^−1^ concentration window (**Figure 5C**). The resulting linear correlation plot resolved an *R*^2^ of 0.96767 (Adj. *R*^2^ = 0.96734). The calculated slope (0.96011 ± 0.01773) and intercept (0.9452 ± 0.30785 pg ⋅ mL^−1^) demonstrate that the non-linear boosting architecture effectively neutralizes the severe down-shift matrix attenuation caused by non-specific interactions within the aromatic pPhR/pPy co-polymer grid.

#### **D.** NfL Verification (WE4 — XGBoost Architecture)

The axonal injury tracking channel (WE4) leveraging the hybrid Apta-MIP protective structure yielded near-perfect chemometric tracking when paired with XGBR, showing a selection *R*^2^ score of 0.99. Out of a massive pool of 9,260 data points, the 100 validation samples plotted across a 0 to 100 pg ⋅ mL^−1^ concentration span reveal close data convergence along the linear ideal line (**Figure 5D**). The model achieved a striking coefficient of determination (*R*^2^ = 0.99581, Adj. *R*^2^ = 0.99577). The calculated slope (0.99299 ± 0.0065) paired with a tight intercept of 0.4232 ± 0.3764 pg ⋅ mL^−1^ confirms that combining the structural selectivity of the Apta-MIP layer with tree-boosting regression models provides superior protection against chemical crosstalk.

## Conclusion

In this work, we presented the design, fabrication, and analytical validation of SMART-NeuroDx a portable, reagent-free, ML-enhanced electrochemical biosensor array engineered for rapid and simultaneous multi-biomarker screening directly in unprocessed biofluids. By combining surface-confined redox-active molecularly imprinted polymers (Sc-RA-MIPs) and Sc-RA-AptaMIP/MIP interfaces on nanostructured porous gold (HPG) PCB electrodes, the platform achieves label-free signal transduction in under 35 minutes. Quantum chemical DFT modelling and molecular docking simulations confirmed the thermodynamic stability of the customized polymer backbones (pPhR, pPy, and PyPEG), while potential-assisted electrostatic gating eliminated cross-channel template contamination during sequential electropolymerization.

To deconvolute complex Faradaic signal responses, a chemometric machine learning pipeline was established. Extracting five multi-dimensional voltammetric features (Δ*I_p_*, Signal Loss %, Δ*E*, ΔAUC, ΔFWHM) and coupling them with 5-fold cross-validated Random Forest and XGBoost regressors mathematically isolated authentic target binding from matrix interferences and crosstalk (*R*^2^ = 0.94-0.99). When interfaced with our custom, STM32G4-based handheld potentiostat, the platform demonstrated benchtop-equivalent performance against commercial workstations in a deployment-ready format. Analytical testing verified sub-picogram sensitivity for pTau217 (LOD = 0.087 pg ⋅ mL^−1^), broad dynamic ranges across all channels (GFAP ≤ 600 pg ⋅ mL^−1^), high operational precision (RSD ≤ 5.34%), and minimal biofouling attenuation (deviation < ±10%) in human plasma and serum.

Building upon the analytical validation, surface characterization, and hardware architecture established in this precursor study, upcoming work will apply the SMART-NeuroDx platform directly to human clinical cohorts to address a major bottleneck in neurodegenerative diagnostics, notably, the cost, infrastructure and the time required for a plasma biomarker panel.

Ultimately, this foundational platform establishes the technological framework for a new class of ML-driven, point-of-care multiplex biosensors capable of transforming single-marker diagnostic workflows into comprehensive, multi-dimensional differential screening systems.

## ASSOCIATED CONTENT

### Supporting Information

The Supporting Information is available free of charge at [ACS Sensors].

Molecular modeling, electron-density, and electrostatic-potential mapping of the PyPEG crosslinker; representative baseline and target-bound differential pulse voltammograms for the four channels; and comparative baseline voltammograms of the custom handheld potentiostat versus a CHI1030A workstation (PDF).

## Acknowledgements

The authors acknowledge the MRC AMED /UK Japan ([MR/X02153X/1](https://www.amed.go.jp/en/news/program/2001C_00016.html)) research grant for funding a Postdoctoral Research position, a Research Assistant position, travel and subsistence and consumables related to these studies. The authors also acknowledge the use of Materials Innovation Factory (MIF), University of Liverpool facilities for conducting the Raman and FTIR ATR measurements. The Authors acknowledge use of Liverpool University Biobank (RRID:SCR_027601) at the University of Liverpool. Whole blood samples were obtained under the research ethics approval number ethics 22/EE/0230.

## CRediT authorship contribution statement

Sudhaunsh Deshpande: Conceptualization, Investigation, Formal analysis, Software, Data curation, Validation, Visualization, Writing – original draft. Krzysztof Pawlak: Methodology, Investigation, Formal analysis. Ajith Mohan Arjun: Conceptualization, Methodology, writing & review. Moulya Prathap – Investigation and logistical support. Cara Evans – Investigation and logistical support. Theresa Al Alam – Investigation and logistical support. Anu Mary Joy - Investigation and logistical support. Sanjiv Sharma: Conceptualization, Methodology, Resources, Supervision, Project administration, Funding acquisition, Writing – original draft, Writing – review & editing.

## Data availability

The data that support the findings of this study are available from the corresponding author upon reasonable request.

