## Supplementary Information for "SMART-NeuroDx: A Reagent-Free Multianalyte Biosensor Platform with Machine-Learning Readout for Point-of-Care Dementia Screening"

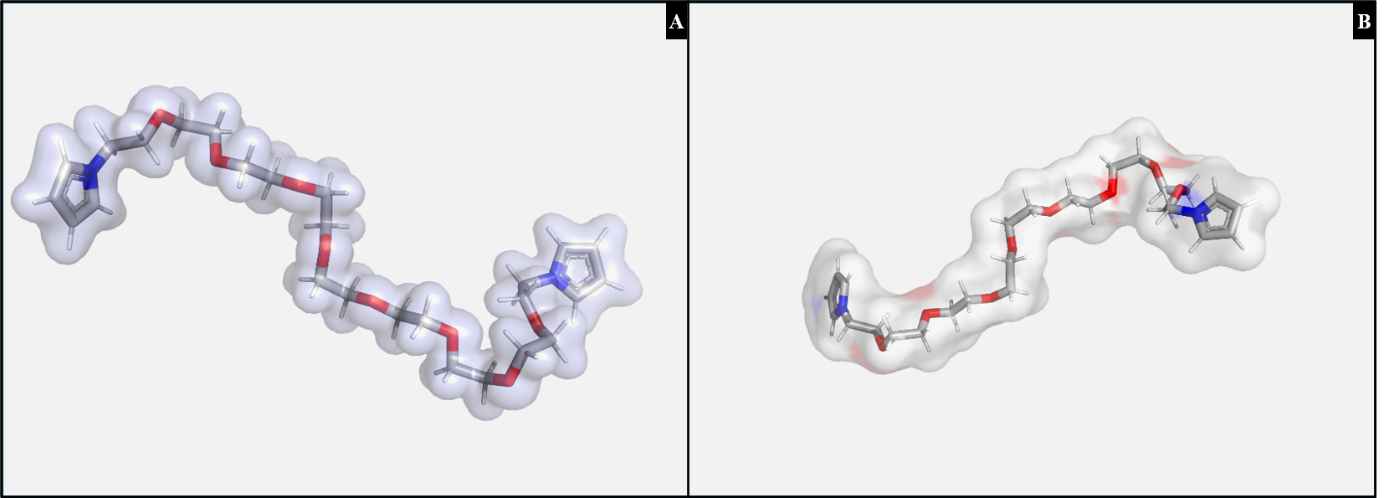

**Figure S1. Molecular modeling, spatial electron density, and electrostatic potential (ESP) mapping of the optimized bis-pyrrole-PEG500 (PyPEG) crosslinker.**

***(A) Quantum mechanical electron density envelope*** *(*$\rho\left( \boldsymbol{r} \right)=0.02\text{ a.u.}$*) enclosing the energy-minimized molecular conformation, illustrating the continuous, high-entropy electron cloud sheathing the flexible PEG backbone and the sterically unhindered, planar terminal pyrrole rings.*

***(B) Electrostatic potential (ESP) mapped onto the solvent-accessible surface (SAS)****, highlighting the localized, highly electronegative (red) pockets over the ether oxygen atoms which coordinate the protective hydration shell, alongside the neutrally balanced (grey/blue) aliphatic domains.*

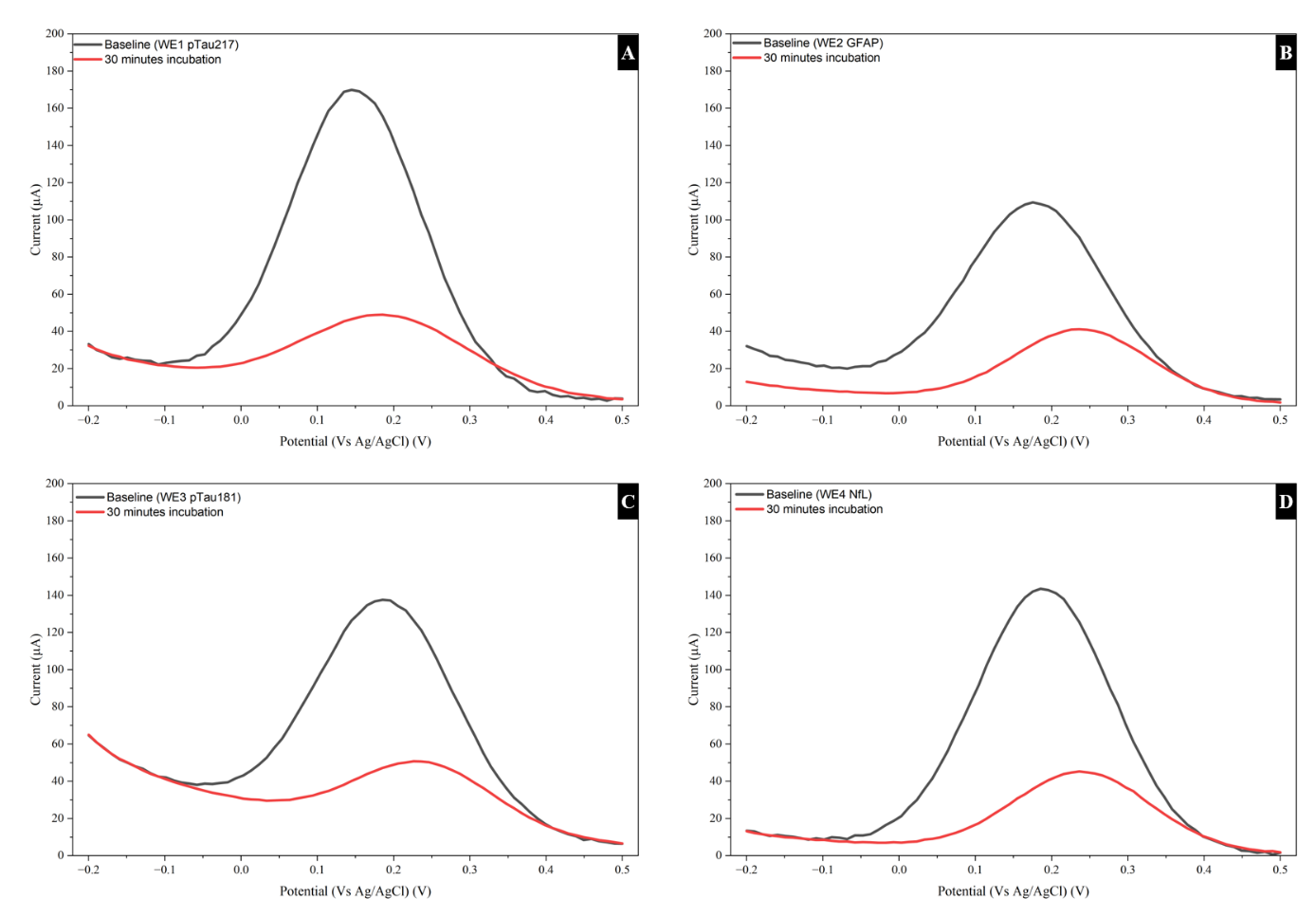

**Figure S2. Representative baseline and target-bound differential pulse voltammetry (DPV) responses across the four biosensor channels.**

Overlaid DPV curves swept from $-0.2\text{ to }+0.5\text{ V}$ (vs. $\text{Ag/AgCl}$) illustrating the intrinsic Faradaic oxidation peak current of the polymer backbones before (black traces) and after a 30-minute incubation with target analytes (red traces):

***(A)*** *WE1 (*$\text{pTau217}$ *channel,* $\text{pPhR/PyPEG}$ *matrix);*

***(B)*** *WE2 (*$\text{GFAP}$ *channel,* $\text{pPhR}$ *matrix);*

***(C)*** *WE3 (*$\text{pTau181}$ *channel,* $\text{pPhR/pPy}$ *matrix); and*

***(D)*** *WE4 (*$\text{NfL}$ *channel,* $\text{AptaMIP}$ *matrix).*

The clear attenuation of the Faradaic oxidation peak current across all channels demonstrates clean label-free signal transduction driven by target binding within the synthetic recognition cavities.

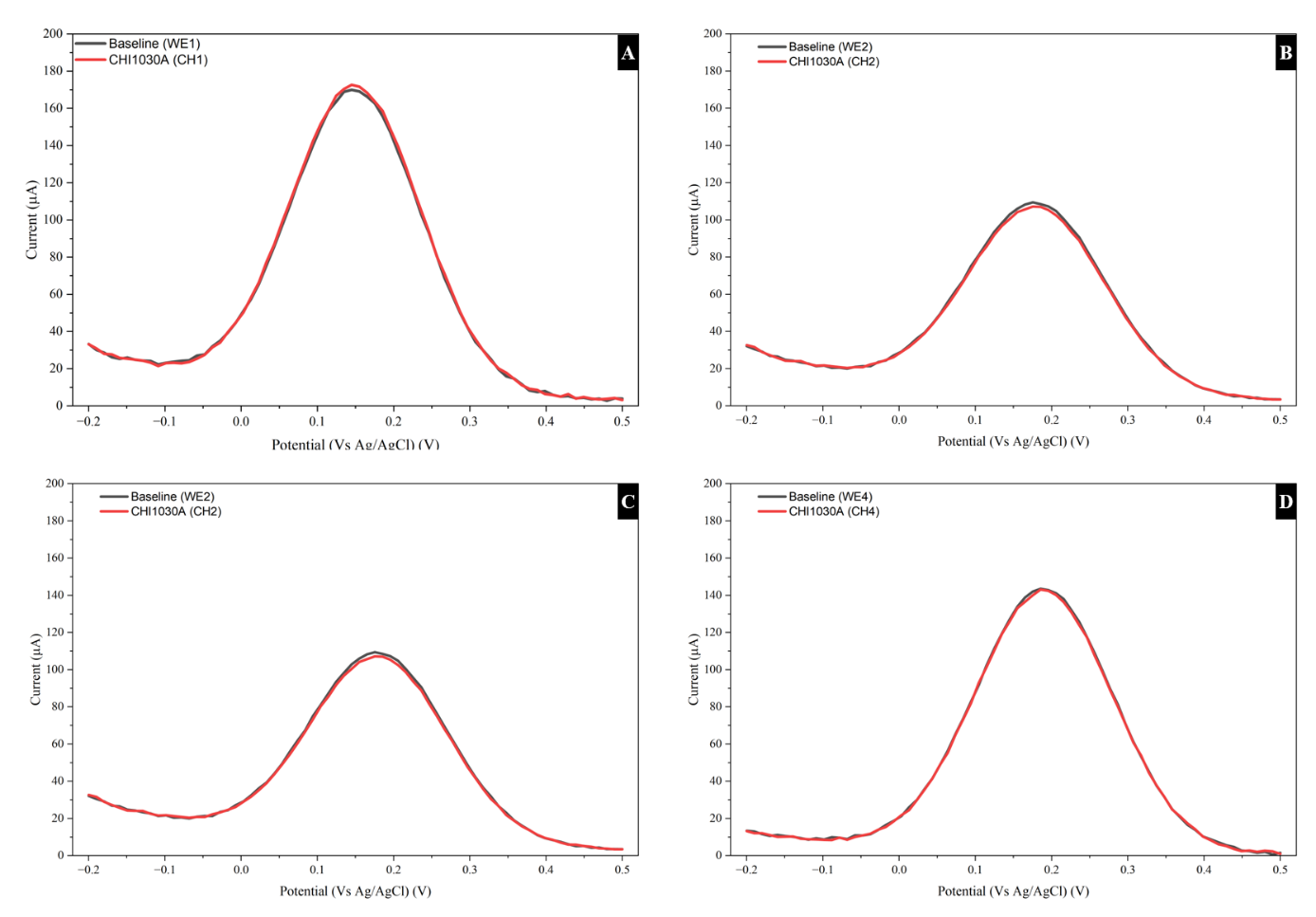

**Figure S3. Comparative baseline DPV voltammograms of the custom potentiostat versus a commercial CHI1030A workstation.**

Overlaid differential pulse voltammetry (DPV) curves recorded in PBS ($-\text{0.2 to }+0.5\text{ V}$ vs. $\text{Ag/AgCl}$) comparing the custom handheld potentiostat (black line) with the CHI1030A multi-potentiostat (red line) across all four working channels:

***(A)*** *Channel 1 (*$\text{WE1}$*, pTau217 channel (pPhR/PyPEG));*

***(B)*** *Channel 2 (*$\text{WE}\text{2, }$ *GFAP channel (pPhR));*

***(C)*** *Channel 3 (WE3, pTau181 channel(pPhR/pPy)); and*

***(D)*** *Channel 4 (WE4, NfL channel(MN711-pPhR/pPy)).*

Supplementary Methods and Results

Computational molecular modelling

To theoretically validate the affinity and selectivity of the synthetic functional monomer configurations prior to electropolymerization, automated molecular docking simulations were performed to evaluate the binding energy interactions between the targeted protein templates and the monomer species. The three-dimensional structures of the target biomarkers—phosphorylated Tau217, Glial Fibrillary Acidic Protein (GFAP), pTau181, and Neurofilament Light chain (NfL) were retrieved from the UniProt ^1^. The spatial structures of the functional monomers, including phenol red, pyrrole, and the synthesized pyrrole-polyethylene glycol (PyPEG) conjugate, were geometrically optimized and energy-minimized using computational density functional theory (DFT) methods ^2^.

Molecular docking simulations were executed using the AutoDock Vina software environment to calculate the lowest-energy binding conformations and free energy of binding ($\Delta G$, $\text{kcal}\cdot\text{mol}^{-1}$) for each monomer-template pair ^3,4^. Prior to docking, all receptor proteins and monomer ligands were converted to. pdbqt format using AutoDockTools by adding polar hydrogen atoms, merging non-polar hydrogens, and assigning Gasteiger partial charges. Grid boxes were parameterized to encompass either the entire protein surface (blind docking for globally distributed cavity analysis) or targeted high-affinity functional domains, such as the specific binding loop of the MN711 aptamer-hybrid interface on WE4. The docking simulations were executed with an exhaustiveness setting of 32 to ensure thorough conformational sampling. The resulting binding energy matrices were analysed to correlate theoretical thermodynamic stability with the empirical sensitivity and crosstalk mitigation observed during multi-channel analytical validation.

Surface Characterization Protocols

Fourier Transform Infrared Spectroscopy (FTIR-ATR): Chemical functional groups and composite matrix compositions were evaluated using a Bruker Vertex 70 spectrometer equipped with a Diamond MIRacle ATR accessory and a DLaTGS detector. Spectra were collected over $4000-400\text{ cm}^{-1}$ with a resolution of $4\text{ cm}^{-1}$ by averaging 32 scans.

Raman Mapping: The spatial distribution and uniformity of the pPhR, pPy, and blended PyPEG matrices across the HPG boundaries were monitored using a Renishaw InVia Raman microscope equipped with a 532 nm laser line (10% power, 1 s exposure). Large-area surface tracking was completed via StreamHR mode using a 10× objective, and polymer components were identified using Renishaw's empty modelling algorithms.

Custom Handheld Potentiostat Hardware and Firmware

To transition the four-channel diagnostic assay into a decentralized Point-of-Care (PoC) platform, a custom, battery-powered handheld potentiostat was designed and developed.

The hardware architecture is cantered on the STM32G491RCT6 microcontroller (STMicroelectronics), featuring a high-performance 32-bit ARM Cortex-M4 core with a floating-point unit (FPU) operating at a clock speed of up to 170 MHz. This device coordinates all waveform generation, high-speed multi-channel data acquisition, peripheral communication, and power optimization routines.

**Real-Time Data Visualization:** User interaction and real-time diagnostic readouts are managed via a high-contrast SSD1309-driven 2.4-inch OLED display (128 × 64-pixel resolution). The display is interfaced using a high-speed I2C communication protocol operating at Fast Mode Plus speeds (up to 1 Mbit/s) to ensure fluid, lag-free UI updates of real-time sweep progress and final concentration outputs.

**Local High-Capacity Data Logging:** To guarantee seamless operation in field settings with unstable remote connections, a microSD card slot is integrated. It interfaces with the STM32 via a dedicated Serial Peripheral Interface (SPI) bus, allowing local, non-volatile data storage of raw time-current transient curves, calibration parameters, and timestamped testing records.

The analogue front-end (AFE) of the device was designed to support robust, high-accuracy electrochemical sweep generation and signal acquisition across widely varying biomarker concentrations:

**Integrated Multi-Stage Electrostatic Discharge (ESD) Protection:** To shield the sensitive, high-impedance internal analogue blocks of the STM32G491RCT6 from physical handling and static discharges during sensor strip insertion, the electrode interface traces (WE, CE, RE) feature low-capacitance ESD protection arrays. These arrays suppress high-voltage transients while keeping leakage currents well below 1 pA to preserve the fidelity of sub-nA Faradaic measurements.

**Hardware Auto-Ranging Circuitry:** Due to the wide concentration variation across the four target biomarkers (ranging from trace levels of pTau to elevated levels of GFAP), the transimpedance amplifier (TIA) feedback network utilizes an active auto-ranging scheme. An array of low-leakage analog switches dynamically shifts the feedback resistors ($R_{f}=10\text{ k}\Omega\text{ to }100\text{ K}\Omega$), seamlessly adjusting the current measurement range on-the-fly. This ensures maximum signal-to-noise ratio (SNR) for picoampere-level currents without saturating the ADC during high-current sweeps.

**Waveform Generation and Signal Conversion:** Excitation sweep profiles are generated using the MCU’s internal, buffered 12-bit Digital-to-Analog Converters (DACs). The resulting current response is routed through the TIA stage and sampled using the integrated 12-bit Analog-to-Digital Converters (ADCs). Real-time hardware oversampling is activated within the MCU to achieve an effective resolution of 16-bit precision, filtering out high-frequency noise directly at the physical layer.

Molecular modelling and Density functional theory:

**Density function theory (DFT):** To establish a high-fidelity structural and thermodynamic baseline for the anti-biofouling PEG matrix, geometry optimization of the synthesized bis-pyrrole-poly (ethylene glycol) (bis-pyrrole-PEG500; $\text{C}_{24}\text{H}_{46}\text{O}_{12}\text{N}_{2}$) crosslinker was executed. To accurately represent the experimental parameters of the poly (ethylene glycol) diglycidyl ether (PEGDE) precursor ($M_{\text{w}}\approx500 \text{g}\cdot\text{mol}^{-1}$), a symmetrical model featuring two terminal pyrrole rings bridged by a 9-unit repeating ethylene glycol chain (80 atoms total) was constructed. The initial geometry was pre-relaxed and subsequently subjected to full quantum mechanical geometry optimization utilizing the GeometRIC solver coupled with DFT calculations at the B3LYP/6-31G(d) level of theory.

The system converged successfully after 44 Self-Consistent Field (SCF) optimization cycles, yielding a stable, global minimum ground-state electronic energy of $-1728.36249249\text{ Hartree}$.

The optimized coordinates reveal a highly stable, folded configuration where the central polyether oxygen atoms coordinate self-stabilizing intramolecular interactions, while leaving the terminal planar pyrrole rings sterically unhindered. This spatial layout is highly advantageous; the unconstrained terminal pyrrole rings remain fully accessible for electropolymerization, enabling the formation of a robust, highly crosslinked network directly on the highly porous gold (HPG) surface finish. Concurrently, the fully relaxed, highly flexible PEG domain is free to extend into the liquid interface to maximize hydration, providing the physical basis for the exceptional anti-biofouling performance and quasi-size-exclusion properties verified on the clinical multi-analyte array.

The fully relaxed, three-dimensional spatial conformation of the optimized bis-pyrrole-PEG500 molecule exhibits several key structural characteristics that directly support its high-performance sensing capability:

**Hydrophilic Oxygen Exposure:** The ether oxygen atoms (represented in red) are highly exposed along the outer envelope of the wavy backbone. When grafted onto the HPG interface, these exposed, highly electronegative oxygen sites are ideally positioned to coordinate water molecules. This hydration shell generation is the fundamental mechanism behind the matrix's exceptional anti-biofouling performance, effectively preventing the non-specific adsorption of bulky serum lipids and proteins while maintaining excellent Faradaic signal-to-noise ratios.

To provide a molecular-level understanding of the physical mechanisms governing the anti-biofouling performance and rapid polymerization kinetics of the synthesized bis-pyrrole-poly (ethylene glycol) ($\text{PyPEG}$) crosslinker, quantum chemical calculations were coupled with spatial electron density and electrostatic potential (ESP) mapping. The optimized ground-state geometry, resolved at the B3LYP/6-31G(d) level of theory, was translated into 3D spatial grids to evaluate the quantum mechanical electron distribution and surface charge characteristics.

**Quantum Mechanical Electron Density Envelope (**$\boldsymbol{\rho}\left( \mathbf{r} \right)$**): Figure S1A** displays the total quantum mechanical electron density mapped as a translucent, lavender-blue volumetric envelope ($\rho\left( r \right)=0.02\text{ a.u.}$) enclosing the energy-minimized stick framework of the $\text{PyPEG}$ molecule:

**Backbone Entropy and Conformational Slack:** The continuous, sinusoidal electron density envelope sheathing the repeating polyether backbone ($-\text{CH}_{2}\text{-CH}_{2}\text{-O-}$) reveals a highly flexible, strain-free wavy conformation. The smooth distribution of the electron cloud over the aliphatic segments highlights the high-entropy nature of the central polymer linker. This structural flexibility allows the chain to adapt dynamically at the liquid-solid interface.

**Terminal Anchor Availability:** At the molecular termini, the electron density over the five-membered pyrrole rings shows clean, planar $\pi$-conjugation. The delocalized $\pi$-electron cloud of the pyrrole rings remains entirely unhindered by the central polyether folding. This ensures that the terminal radical-forming centers are fully accessible for unconstrained $\pi$-$\pi$ stacking and low-defect electropolymerization directly onto the highly porous gold (HPG) working electrode

**Electrostatic Potential (ESP) Surface: Figure S1B** displays the electrostatic potential mapped onto the solvent-accessible surface (SAS) of the monomer, mathematically illustrating the charge boundaries that dictate interfacial water coordination:

**Hydrophilic Oxygen Pockets (Red Regions):** The ESP map reveals localized, highly electronegative (electron-rich, red) pockets concentrated directly over the ether oxygen ($\text{O}$) atoms along the inner curves of the undulating polyether backbone. These concentrated negative potential zones act as strong nucleophilic coordinating centres. When the $\text{PyPEG}$ matrix is grafted onto the electrode surface, these red oxygen sites actively coordinate and secure a dense, rigid network of hydrogen-bonded water molecules, generating the physical hydration shell that repels non-specific biofouling.

**Electroneutrally Balanced Sheath (Grey/Blue Regions):** The aliphatic carbon-hydrogen ($\text{C-H}$) segments and the terminal pyrrole rings exhibit predominantly neutral-to-weakly positive (grey-to-blue) electrostatic potentials. This balanced charge distribution prevents native self-aggregation or electrostatic collapse of the polymer matrix, ensuring that the embedded molecularly imprinted polymer (MIP) cavities remain stable, open, and fully accessible to target biomarkers.

To theoretically justify the monomer compositions chosen for each biomarker channel ($\text{WE1 - WE4}$), automated molecular docking simulations were conducted via AutoDock Vina ^3,4^. The thermodynamic affinity—expressed as the lowest free energy of binding ($\Delta G$, $\text{kcal}\cdot\text{mol}^{-1}$)—was systematically mapped for each targeted dementia biomarker against three distinct receptor matrix candidate environments: pure $\text{pPhR}$, a blended $\text{pPhR/pPy}$ composite, and a $\text{pPhR/PyPEG}$ matrix. The calculated binding energy profiles are summarized in **Table S1**.

**Table S1: AutoDock Vina Binding Affinity Matrix and Monomer Optimization Profiles**

| Target Biomarker | pPhR (kcal⋅mol−1) | pPhR/pPy (kcal⋅mol−1) | pPhR/PyPEG (kcal⋅mol−1) | Chosen Matrix Structure | Functional Justification |
| --- | --- | --- | --- | --- | --- |
| **pTau217** | -3.92 | -5.197 | -8.62 | $\text{pPhR/PyPEG}$ | Ultra sensitivity |
| **GFAP** | -5.63 | -8.63 | -7.82 | $\text{pPhR}$ | Extended Range |
| **pTau181** | -4.67 | -13.10 | -6.52 | $\text{pPhR/pPy}$ | Ultra sensitivity |
| **NfL** | -5.20 | -8.30 | -6.20 | $\text{pPhR/pPy}$ | Ultra sensitivity |

Thermodynamic Justification of Matrix Allocation

For the pTau217 screening channel, the $\text{pPhR/PyPEG}$ blended matrix yielded the highest thermodynamic binding affinity with a pronounced $\Delta G$ of $-8.62 \text{kcal}\cdot\text{mol}^{-1}$, significantly outperforming the pure $\text{pPhR}$ ($-3.92 \text{kcal}\cdot\text{mol}^{-1}$) and $\text{pPhR/pPy}$ ($-5.197 \text{kcal}\cdot\text{mol}^{-1}$) configurations. This strong interaction energy stems from the favorable spatial orientation of the template within the oxygen-rich polyether loops of the optimized PEG framework. Consequently, this configuration was chosen to achieve the sub-picogram ultra sensitivity ($0.087 \text{pg}\cdot\text{mL}^{-1}$) required to detect trace pTau217 in early-stage clinical plasma samples.

In contrast to the tauopathy markers, the selection criteria for the Glial Fibrillary Acidic Protein (GFAP) channel prioritized an extended dynamic range over extreme trace sensitivity, matching its elevated concentrations during active astrocytic neuroinflammation. Although the $\text{pPhR/pPy}$ matrix exhibited the tightest binding energy ($-8.63 \text{kcal}\cdot\text{mol}^{-1}$), the pure $\text{pPhR}$ matrix was strategically chosen. Its moderate, well-distributed docking energy of $-5.63 \text{kcal}\cdot\text{mol}^{-1}$ avoids rapid site saturation, which successfully extended the analytical working scope up to $600 \text{pg}\cdot\text{mL}^{-1}$ while preventing early plateauing of the Faradaic signal response.

The $\text{pPhR/pPy}$ composite matrix displayed an exceptionally strong affinity for both pTau181 ($-13.100 \text{kcal}\cdot\text{mol}^{-1}$) and NfL ($-8.300 \text{kcal}\cdot\text{mol}^{-1}$). The high electronic density and structural rigidity of the co-polymerized pyrrole rings provide robust non-covalent $\pi$-$\pi$ stacking and electrostatic coordination sites that securely lock the template geometries. This strong binding profile was exploited for both WE3 and WE4 to achieve the ultra-sensitivity profiles required for high-fidelity detection of classic tauopathy and general axonal degeneration thresholds.

Fourier Transform Infrared Attenuated Total Reflectance

To confirm the functional group identities, copolymerization pathways, and crosslinker integration across the independent channels, Fourier-Transform Infrared Attenuated Total Reflection (FTIR-ATR) spectroscopy was conducted over the $400\text{ to }4500\text{ cm}^{-1}$ wavenumber range. The independent channel spectra provide clear chemical signatures distinguishing the uniquely functionalized surface matrices.

FTIR-ATR Spectroscopic Characterization

**WE1 (pPhR/PyPEG):** The fingerprint region (700–1500 cm⁻¹) shows C–H bending (~790 cm⁻¹), C=C bending (~1000 cm⁻¹), C–O phenol stretching (~1380 cm⁻¹), and a diagnostic aliphatic ether C–O stretch (~1120 cm⁻¹) confirming PEG spacer incorporation. Conjugated C=O stretching appears at ~1680 cm⁻¹, with aliphatic C–H (~2850 cm⁻¹) and pyrrole-derived N–H stretching (~2930 cm⁻¹) further upfield. A broad, intense O–H stretch centered at ~3400 cm⁻¹ (3100–3650 cm⁻¹) reflects the hydrated, hydrogen-bonding-active PEGylated matrix (**Figure S5A**).

**WE2 (pPhR):** Dominated by sulfonate/phenolic features: C–H bending (790 cm⁻¹), S=O stretching (1030 cm⁻¹), S=O stretching in phenol (1250 cm⁻¹), O–H bending in phenol (1360 cm⁻¹), C–H bending (1470 cm⁻¹), and a sharp quinoid C=O stretch at 1680 cm⁻¹. A shallow aliphatic C–H stretch (~2920 cm⁻¹) sits beneath a broad O–H stretch (3050–3680 cm⁻¹) from the unmodified hydroxyl network (**Figure S5B**).

**WE3/WE4 (pPhR/pPy):** Near-identical replicate profiles confirm reproducible co-deposition. Key bands: S=O stretching (1030 cm⁻¹, residual PhR), C–O stretching (1140 cm⁻¹), C–N stretching (1260 cm⁻¹), S=O bending (1350 cm⁻¹), C–H bending (1460 cm⁻¹), N–H bending (1620 cm⁻¹), and conjugated ring C=C stretch (1680 cm⁻¹). A small S–H stretch (~2600 cm⁻¹) indicates thiol-anchoring functionality, alongside aliphatic C–H (2830 cm⁻¹) and pyrrole N–H stretching (2930 cm⁻¹). The O–H stretch band (3200–3650 cm⁻¹) is weaker than in WE1, consistent with lower PEG-derived hydration (**Figure S5C**).

**Comparative Overlay** (**Figure S5D**): All four electrodes preserve the core aromatic skeleton (1200–1700 cm⁻¹). WE1 shows distinct suppression and smoothing at the C–O–C ether stretch (1120 cm⁻¹) and O–H hydration region (3400 cm⁻¹), confirming successful PyPEG intercalation into the matrix.

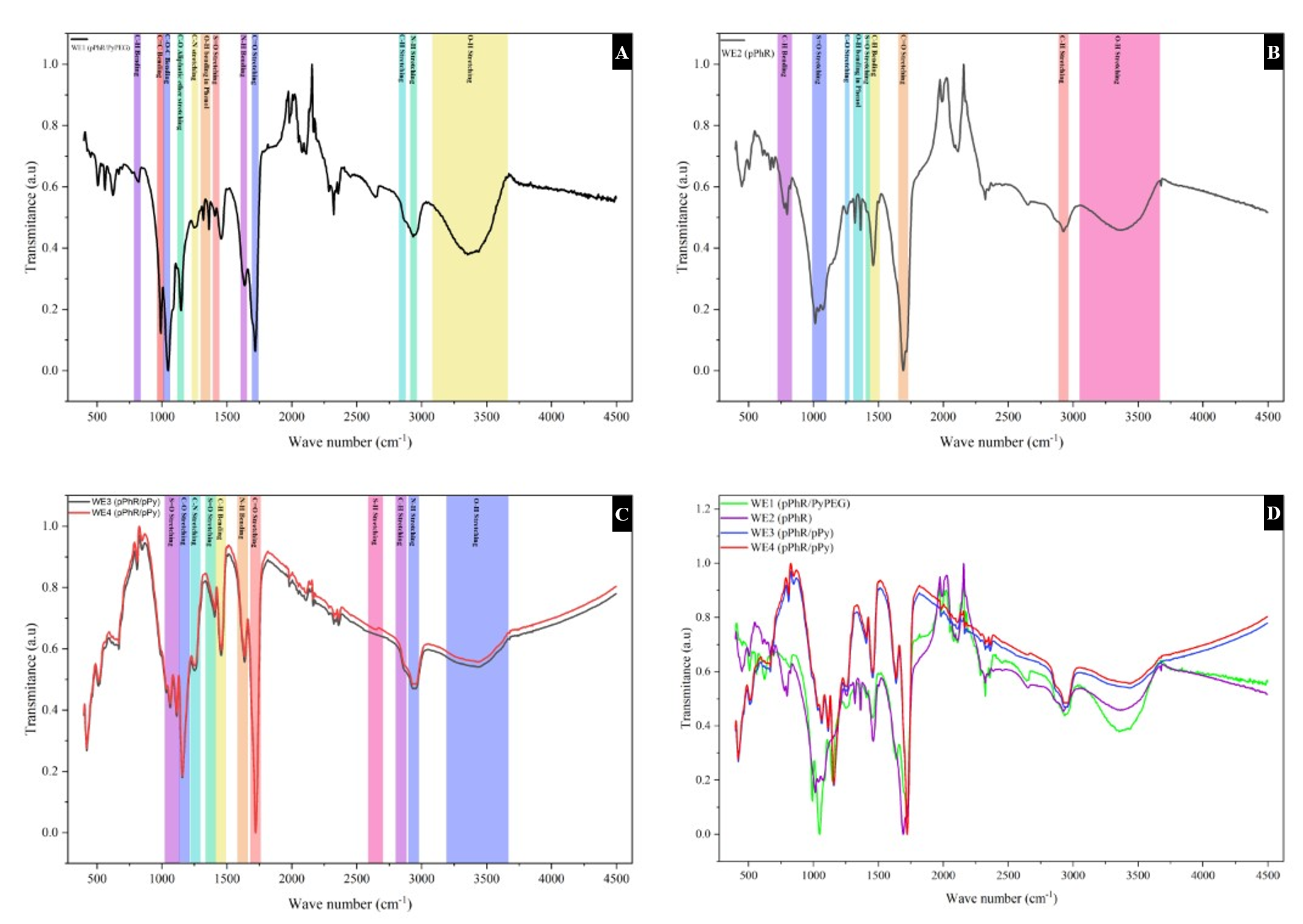

**Figure S4. FTIR-ATR spectra of the functionalized sensing matrices across the four-channel array.**

Individual baseline-corrected transmission spectra are shown with color-coded vibrational band assignments for:

**(A) WE1** functionalized with the pegylated ternary $\text{pPhR/PyPEG}$ crosslinked matrix.

**(B) WE2** functionalized with the pure $\text{pPhR}$ homopolymer matrix.

**(C) WE3** (black) **WE4** (red) functionalized with identical binary $\text{pPhR/pPy}$ composite matrices, detailing high co-deposition alignment.

**(D)** Comparative spectral overlay across all four working channels from $400\text{ to }4500\text{ cm}^{-1}$, highlighting the specific baseline adjustments and ether/hydroxyl stretching enhancements driven by channel-specific functionalization.

Hardware instrumentation of the handheld potentiostat

To transition the multiplexed Sc-RA-MIP sensor strip into a point-of-care system, a custom four-channel handheld potentiostat was built around the STM32G491RCT6 microcontroller. The architecture integrates on-chip analogue peripherals to execute synchronized multi-channel differential pulse voltammetry (DPV) without external signal generators. Crucially, to ensure seamless translation to point-of-care deployment, all multi-channel voltammetric feature extraction and machine learning dataset collection were executed directly on the custom handheld potentiostat, with initial device signal fidelity benchmarked against the CHI1030A workstation.

**Excitation & Reference Feedback Control:** Excitation waveforms generated by an integrated 12-bit DAC pass through a differential driver stage (Diff) to drive the counter electrode (CE) (OPA2140AIDR). Reference potential stability is maintained via a unity-gain buffer amplifier (Buff) connected to the reference electrode (RE) relative to a virtual ground (VGND) (1.65 V to analogue ground (AGND)). Direct Memory Access (DMA) channels stream excitation vectors without CPU overhead (**Figure S5**).

**Transimpedance Amplification (TIA) & Active Auto-Ranging:** Each working electrode channel ($\text{WE1-WE4}$) feeds directly into a dedicated transimpedance amplifier (TIA) (OPA2140AIDR) referenced to VGND (1.65 V to AGND). To accommodate broad dynamic ranges across different polymer matrices, each channel features software-selectable feedback gain resistors ($R_{f}$ and $10R_{f}$) (**Figure S5**).

**Synchronized Acquisition & User Interface:** Analog outputs from the TIAs are routed to an array of high-speed 12-bit ADCs managed by the DMA controller. Real-time measurements are rendered on a 2.4-inch SSD1309 OLED display via an I2C bus and recorded onto a microSD card over a SPI interface (**Figure S5;** actual device shown in **Figure 1**).

Benchtop Validation against CHI1030A

To verify analytical fidelity, baseline DPV scans obtained using the custom handheld potentiostat were benchmarked directly against a commercial CHI1030A laboratory electrochemical workstation across all four channels. As shown in **Figure S3**, the baseline voltammograms generated by the custom device (black traces) track the CHI1030A curves (red traces) across the $-0.2\text{ to }+0.5\text{ V}$ potential range. Peak potential alignments ($\sim+0.15\text{ to }+0.18\text{ V}$) and Faradaic peak current magnitudes are identical across all channels (WE1-WE4), confirming benchtop-grade electrochemical precision in a portable, battery-powered format.

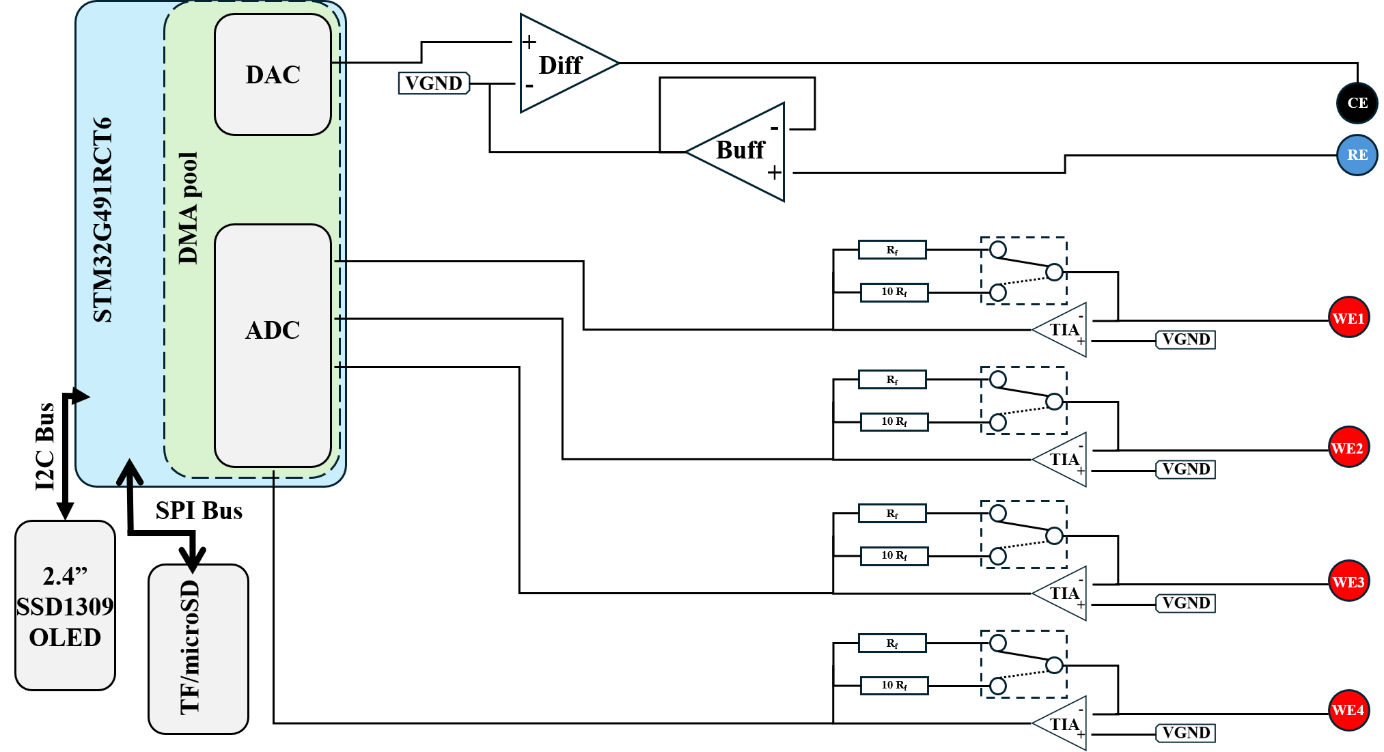

**Figure S5. Architectural block diagram of the custom STM32G4-based four-channel handheld potentiostat.** The embedded system utilizes an STM32G491RCT6 microcontroller featuring a Direct Memory Access (DMA) pool driving on-chip DACs and ADCs. The excitation module comprises a differential driver amplifier (Diff) and reference buffer (Buff) driving the counter (CE) and reference (RE) electrodes relative to a virtual ground (VGND). Current responses from the four independent working electrodes ($\text{WE1-WE4}$) are converted via dedicated transimpedance amplifiers (TIA) with active dual-gain feedback control ($R_{f}$ / $10R_{f}$). Peripherals include an SSD1309 OLED display driven over $\text{I}^{2}\text{C}$ and a microSD card logger communicating via SPI.
